# Enhancing radiotherapy response via intratumoral injection of the TLR9 agonist CpG to stimulate CD8 T cells in an autochthonous mouse model of sarcoma

**DOI:** 10.1101/2024.01.03.573968

**Authors:** Chang Su, Collin L. Kent, Matthew Pierpoint, Warren Floyd, Lixia Luo, Nerissa T. Wiliams, Yan Ma, Brian Peng, Alexander L. Lazarides, Ajay Subramanian, Jonathan E. Himes, Vincent M. Perez, Rosa D. Hernansaiz-Ballesteros, Kimberly E. Roche, Jennifer L. Modliszewski, Sara R. Selitsky, Mari Shinohara, Amy J. Wisdom, Everett J. Moding, Yvonne M. Mowery, David G. Kirsch

**Author notes:** Lead Contact and Corresponding Author: David G. Kirsch, 610 University Ave Toronto, Ontario M5G 2M9, Canada. Chang Su and Collin Kent are co-first authors.

## Abstract

Radiation therapy is frequently used to treat cancers including soft tissue sarcomas. Prior studies established that the toll-like receptor 9 (TLR9) agonist cytosine-phosphate-guanine oligodeoxynucleotide (CpG) enhances the response to radiation therapy (RT) in transplanted tumors, but the mechanism(s) remain unclear. Here, we used CRISPR/Cas9 and the chemical carcinogen 3-methylcholanthrene (MCA) to generate autochthonous soft tissue sarcomas with high tumor mutation burden. Treatment with a single fraction of 20 Gy RT and two doses of CpG significantly enhanced tumor response, which was abrogated by genetic or immunodepletion of CD8+ T cells. To characterize the immune response to RT + CpG, we performed bulk RNA-seq, single-cell RNA-seq, and mass cytometry. Sarcomas treated with 20 Gy and CpG demonstrated increased CD8 T cells expressing markers associated with activation and proliferation, such as Granzyme B, Ki-67, and interferon-γ. CpG + RT also upregulated antigen presentation pathways on myeloid cells. Furthermore, in sarcomas treated with CpG + RT, TCR clonality analysis suggests an increase in clonal T-cell dominance. Collectively, these findings demonstrate that RT + CpG significantly delays tumor growth in a CD8 T cell-dependent manner. These results provide a strong rationale for clinical trials evaluating CpG or other TLR9 agonists with RT in patients with soft tissue sarcoma.

## Introduction

Soft tissue sarcomas are a heterogeneous group of malignancies (1). Following local therapies with surgery and radiotherapy, approximately 50% of patients with large, high-grade tumors develop distant metastasis. After metastases occur, limited therapeutic options are available with median survival of approximately 12 to 18 months (2). Thus, there is a pressing need for alternative therapeutic approaches to improve overall survival for these patients.

Immunotherapy has recently emerged as a promising treatment for many solid tumors (3), with particularly high response rates in melanoma and non-small cell lung carcinoma (NSCLC). However, a phase II clinical trial in advanced soft tissue sarcoma (STS) patients showed that only ∼17.5% of patients responded to anti-PD-1 monotherapy, which suggests that sarcomas are more resistant to immune checkpoint blockade compared to 40-45% objective response rates in melanoma and NSCLC patients (4). Therefore, it is important to explore other forms of immunotherapy that may improve outcomes for patients with sarcoma.

Over 40 years ago, Stone and colleagues used a transplanted model of soft tissue sarcoma to discover that activating the immune system through bacterial infection can enhance tumor control when administered with radiation therapy (RT) (5). Many subsequent studies have suggested that RT can work synergistically with immunotherapy to suppress tumor growth (3, 6, 7). In this study, we found that the combination of CpG (unmethylated cytosine-phosphorothioate-guanosine forms of DNA), a Toll-like receptor 9 (TLR9) agonist, and RT suppresses tumor growth significantly using autochthonous mouse models of soft tissue sarcoma in which the tumor gradually develops under surveillance by an intact immune system (8, 9). Here, we show that CD8^+^ T cells are essential for mediating the anti-tumor effects by CpG and radiotherapy. We further demonstrated that depleting lymphocytes, especially CD8^+^ T cells, negates the treatment effects of CpG. Unlike immune checkpoint inhibitors that aim to reverse the exhaustion state of T cells, the TLR9 agonist CpG combined with RT draws CD8^+^ T cells expressing markers associated with activation and proliferation into the tumor. Taken together, these findings suggest a promising treatment option of combining TLR9 agonists and radiotherapy in treating patients with STS, which often contain few T cells.

## Results

### CpG and radiation therapy suppress autochthonous p53/MCA sarcoma growth

To investigate whether the combination treatment of CpG and RT improves tumor growth delay compared to single therapy with CpG alone or RT alone in a primary tumor model, we induced STS with a high mutational load in 129/SvJae mice by injecting an adenovirus expressing CRISPR-Cas9, a single guide RNA (sgRNA) targeting *Trp53* (sgp53) (8), and the carcinogen 3-methylcholanthrene (MCA) into the gastrocnemius muscle (9) (Figure 1A). After tumor induction, primary sarcomas (p53/MCA model) develop at the injection site over two to three months under the selective pressure of immunoediting in immunocompetent mice (10). p53/MCA sarcomas demonstrate gross and histologic morphologies as well as transcriptional profiles similar to human undifferentiated pleomorphic sarcoma (UPS) (8, 11). When tumor volumes reached 70-150 mm^3^, mice were randomized to receive 0 Gy or 20 Gy RT (Day 0) and CpG or control (GpC dinucleotides with the positions of cytosine and guanine reversed relative to the phosphate linker) (Day 3 and Day 10) (Figure 1B). Significant tumor growth delay was observed with either CpG alone or RT alone compared to the control group. Mice treated with CpG and RT exhibited the longest time to tumor quintupling with a mean of 27.2 days compared to 8.3 days for mice treated with control, 11.9 days for CpG alone, and 20.6 days for radiotherapy alone, suggesting that TLR9 agonist improves radiotherapy’s treatment effect in delaying tumor growth (Figure 1C, 1D).

**Fig 1.**
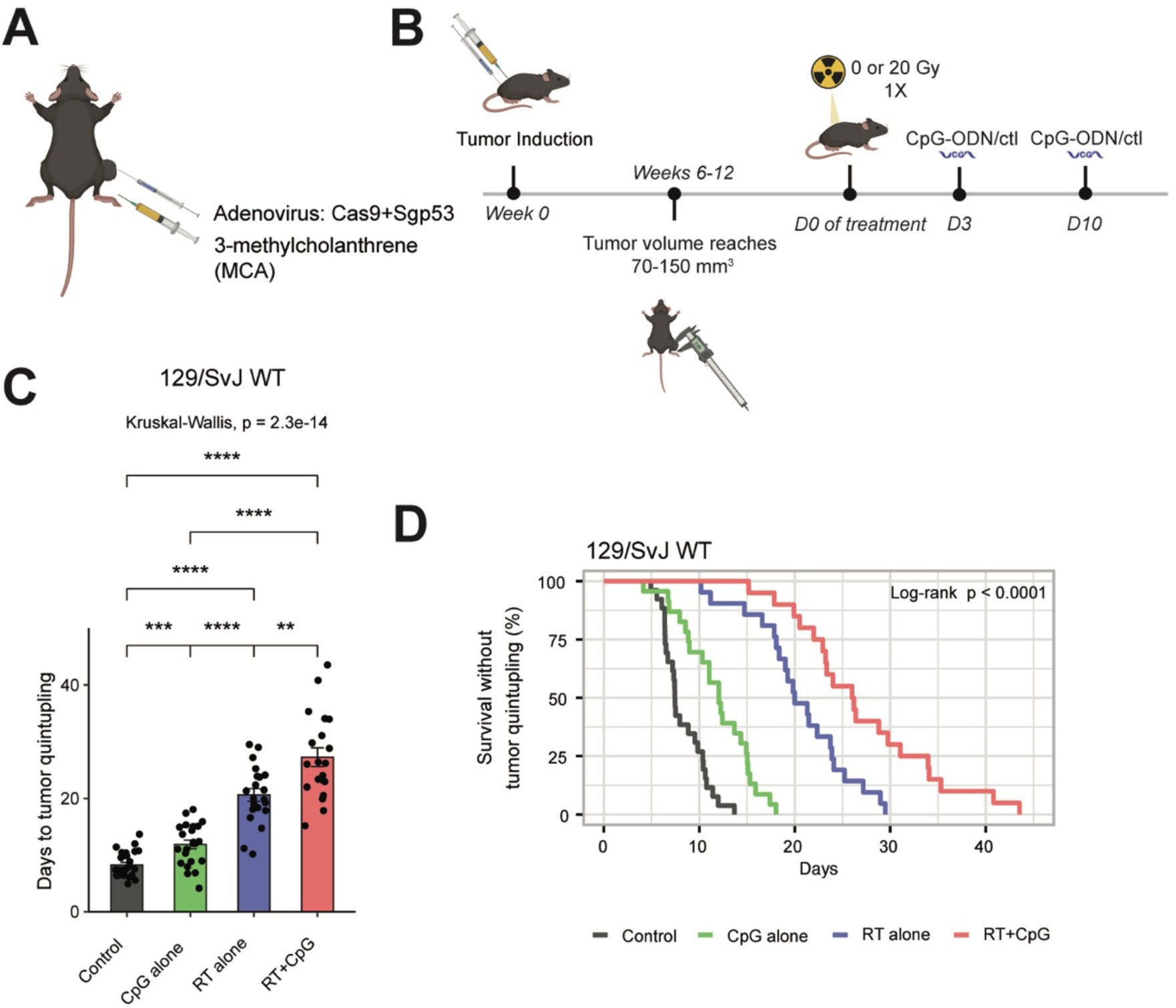
Increased tumor growth delay after treatment with CpG ODN and radiation therapy in autochthonous p53/MCA sarcomas. A. Primary sarcoma initiation by intramuscular injection of Adeno-Cas9-sgp53 and MCA. **B**. Autochthonous sarcomas develop at the injection site about 7-11 weeks after injection. Mice were treated with CpG ODN or control GpC dinucleotides and 0 or 20 Gy when tumors reached >70 mm^3^. **C**. Mice with p53/MCA sarcomas received control GpC dinucleotides with 0 Gy (solid black, *n* = 26), CpG ODN alone (solid green, *n =* 23), control GpC dinucleotides with 20 Gy (solid blue, *n =* 21), or CpG ODN with 20 Gy (solid red, *n =* 20). Time to tumor quintupling (days) after the indicated treatment. **D**. Mice with p53/MCA sarcomas received control GpC dinucleotides with 0 Gy (solid black, *n* = 26), CpG ODN alone (solid green, *n =* 23), control GpC dinucleotides with 20 Gy (solid blue, *n =* 21), or CpG ODN with 20 Gy (solid red, *n =* 20). Kaplan-Meier analysis with tumor quintupling as the endpoint. Kruskal-Wallis Test was used for comparison across the groups, while the Wilcoxon test was selected for the pair-wise comparisons. * shows the significance of the P Value (***: p <= 0.001, ****: p <= 0.0001)

### CyTOF demonstrates significantly increased activated CD8^+^ T cells in CpG + RT treated tumors

To begin to investigate if CpG is mediating growth delay by acting on immune cells rather than killing tumor cells directly, we performed an IncuCyte Live-Cell assay to monitor cell proliferation of three p53/MCA tumor cell lines after co-incubation with titrated concentrations of CpG. The *in vitro* assay demonstrated that CpG does not directly inhibit proliferation of p53/MCA tumor cell lines (Supplementary Figure 1), indicating that *in vivo* tumor growth delay induced by CpG alone is not through direct tumor cell killing. To determine which cell populations play an important role in mediating the treatment effects of CpG and RT, we performed mass cytometry (CyTOF) on tumor cells and tumor-infiltrating immune cells (Figure 2A) (12). We induced p53/MCA tumors as described above and initiated treatment when tumors reached 180-300 mm^3^. We investigated the tumor immune microenvironment on Day 3 after CpG (Day 6 after RT) because we typically observed prominent tumor shrinkage at this time point in mice receiving combination therapy. Figure 2B shows a UMAP plot of CD45 positive cells characterized by CyTOF for the four treatment groups. We observed significantly more CD8^+^ T cells in tumors from mice treated with CpG and RT (Figure 2C) compared to the other treatment groups, which was confirmed by CD8 immunohistochemistry (IHC) staining (Figure 2D, 2E). These CD8^+^ T cells co-expressed Granzyme B and Ki-67, indicating that these CD8^+^ T cells are not exhausted, but rather actively proliferating and primed for cytotoxic cell killing (Figure 2F, G; Supplementary Figure 2).

**Fig 2.**
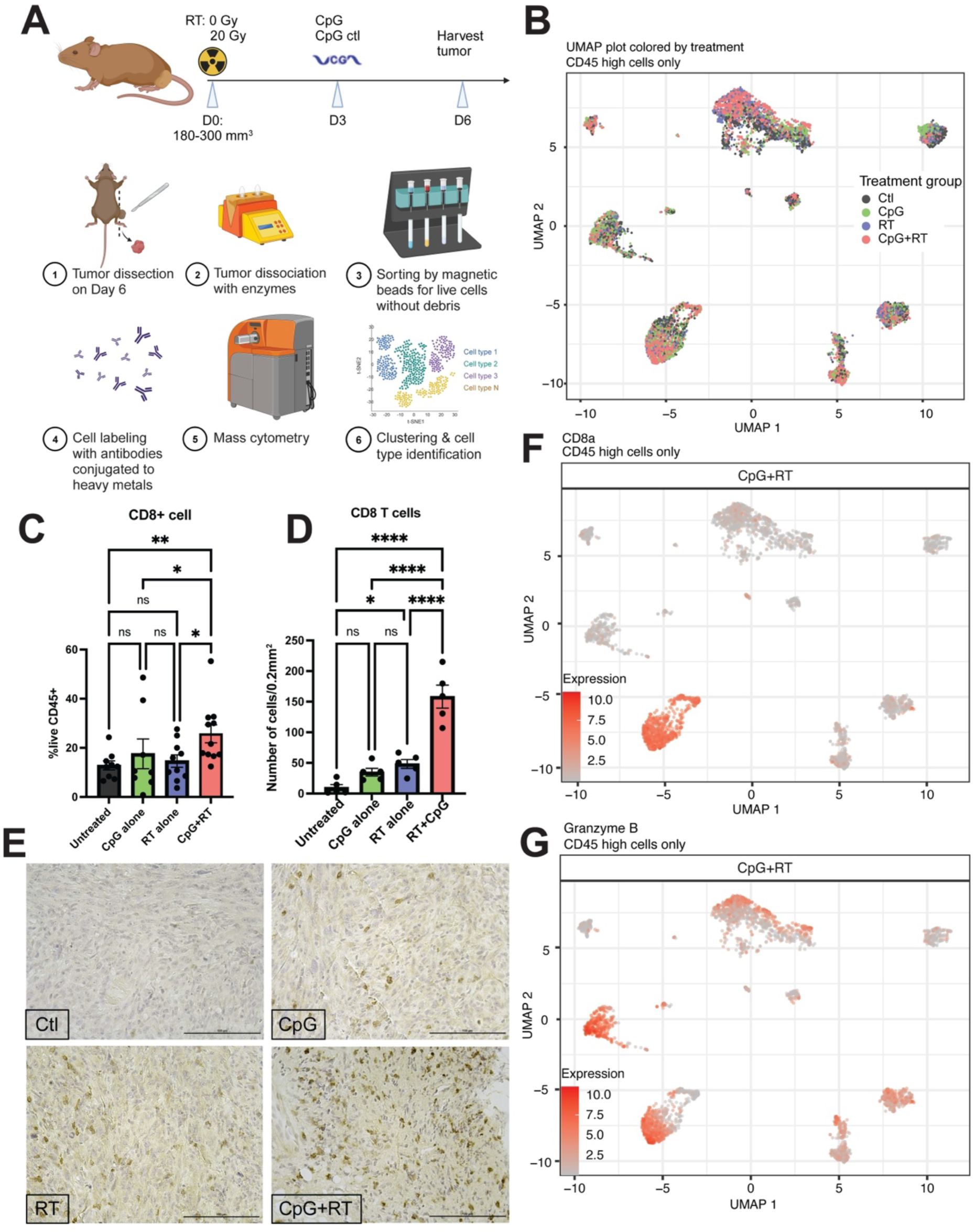
CyTOF and IHC staining demonstrates enhanced intratumoral infiltration of activated CD8 T cells after combination treatment. **A.** Treatment schedule and tumor processing schematic. **B.** UMAP plot of CyTOF data clustering for all CD45 high cells from all tumors and treatment groups. **C.** Frequency of CD8+ cells/live CD45+ cells by CyTOF. Data show mean ± SEM, analyzed by three-way ANOVA. **D.** Average number of CD8+ cells/0.2 mm^2^ in IHC slides. **E.** Representative IHC staining with CD8 Ab. Scale bar = 100 µm. **F.** CD8 expression in CyTOF CD45 high UMAP plot. **G.** Granzyme B expression in CyTOF CD45 high UMAP plot.

### Single-cell RNAseq reveals changes in the adaptive immune system after CpG + RT combination therapy

To identify the major transcriptional differences in immune cell populations with and without CpG and RT, we performed single-cell RNA sequencing (scRNA-seq) on FACS-sorted CD45 positive tumor-infiltrating immune cells from an independent cohort of sarcomas harvested 6 days after 0 or 20 Gy RT and 3 days after Control or CpG (Figure 3A). After filtering and quality control, scRNA-seq analysis generated data for 94,093 cells. Unbiased clustering using shared nearest neighbor (SNN) modularity optimization identified 20 cell clusters with distinct transcriptional profiles that were assigned to known cell lineages utilizing Seurat (13) (Supplementary Figure 3A). The predominant cell populations of intratumoral immune cells were myeloid cells, which is consistent with previous scRNA-seq analyses for primary p53/MCA sarcomas (10).

**Fig 3.**
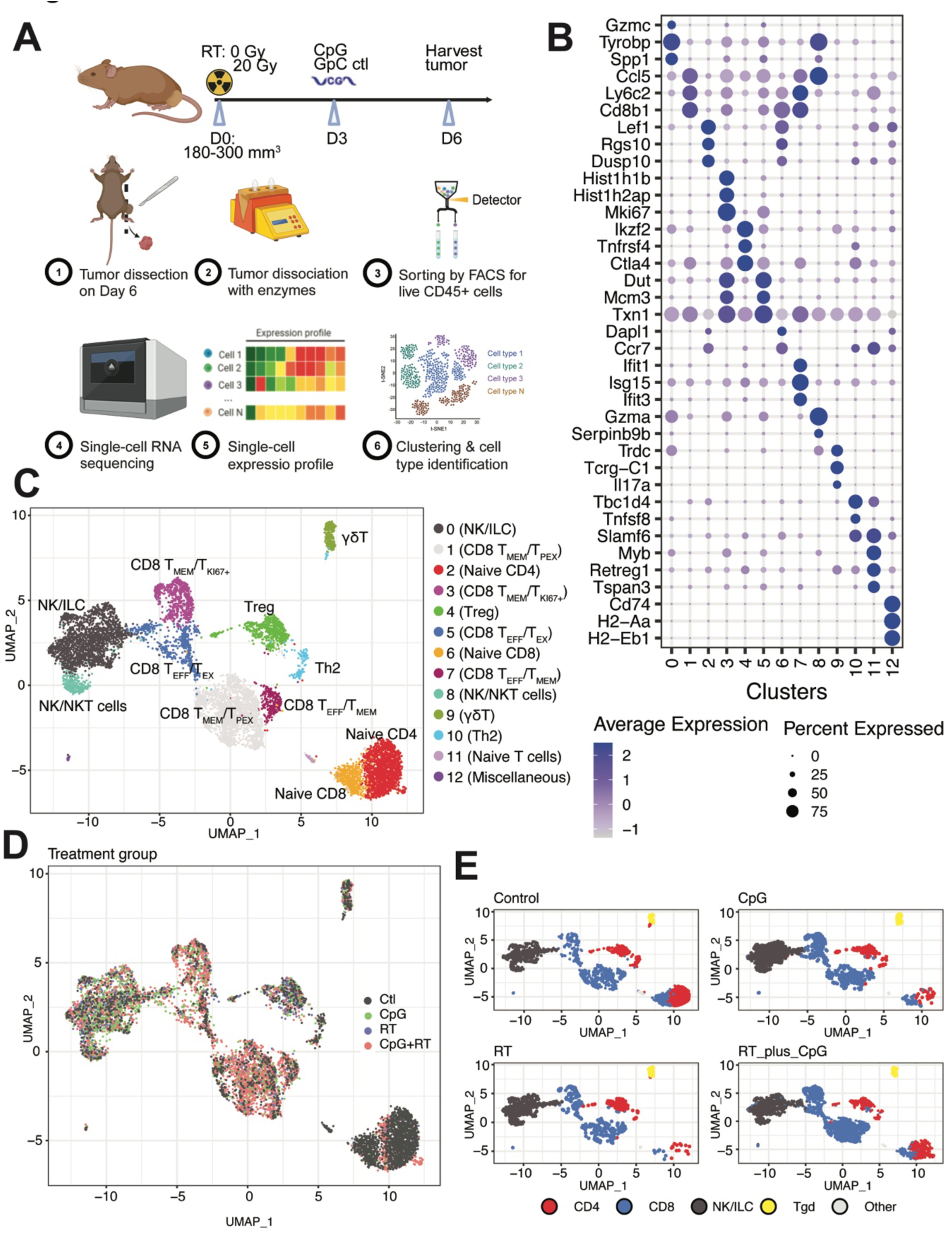
Single cell RNA-seq shows increased CD8 T cell infiltration into the tumor after RT + CpG. **A.** Treatment schedule and tumor processing schematic. **B**. Bubble plot of top 3 differentially expressed genes in each of the T cell subclusters. The shades of color are correlated with levels of expression. The sizes of circles are correlated with percentage of cells in that cluster that express the gene of interest. **C**. UMAP plot of T cells and NK cells scRNA-seq subclustering. **D**. UMAP plot of T cells and NK cells subclustering colored by treatment groups. **E**. UMAP plot of lymphocytes subclustering colored by CD4 (red), CD8 (blue), NK/ILC (black), and γδT (yellow) cells. Mice with p53/MCA sarcomas received control GpC dinucleotides with 0 Gy (*n* = 5), CpG ODN alone (*n =* 5), control GpC dinucleotides with 20 Gy (*n =* 5), or CpG ODN with 20 Gy (*n =* 5).

Since CD8^+^ T cells exhibited the most significant increase in cell numbers based on CyTOF, we sub-clustered T cells and NK cells to better identify transcriptional differences among sarcomas from the four treatment groups. Unsupervised clustering analysis resulted in the identification of 13 subpopulations. The 13 clusters were further annotated based on differentially expressed genes (DEGs) and canonical immune markers, including natural killer cells/innate lymphoid cells/NKT cells (c0 and c8), CD4^+^ T cells (c2, c9, and c10), Treg (c4), and CD8^+^ T cells (c1, c3, c5, c6, and c7) (Figure 3B, 3C, Supplementary Table 1). The most prominent difference across treatment groups was found in the CpG+RT treatment group with a higher number of activated CD8^+^ T cells (c1, c3, c5, c7), which express elevated levels of Granzyme B, Granzyme K, and IFNγ (Figures 3D, 3E Supplementary Figures 3B, 3C, 4, 5). Cell cycle distribution for these clusters is shown in Supplementary Figure 3D, which demonstrates that the majority of the CD8^+^ T cells are actively proliferating and not yet exhausted.

The majority of CD8^+^ T cells in the combination treatment group clustered to c1, c3, c5, and c7. C1 expresses high levels of *Ccl5, Ly6c2, Cxcr6, Cd28, Gzmk, Ifng, Tnfaip3,* and *Tnfaip8* (Figure 3B, Supplementary Figures 3B). *Ccl5* and *Ly6c2* are highly associated with the maintenance and homing of memory T cells (14–16). Connolly and colleagues also identified a group of memory CD8^+^ T cells expressing high levels of *Ccl5* and *Ly6c2* in C57BL/6 mice that were infected with acute lymphocytic choriomeningitis virus (LCMV) (16). *Cxcr6* is a marker for resident memory T cells and antitumor efficacy (17–19). *Cd28, Gzmk, Ifng, Tnfaip3,* and *Tnfaip8* are activation markers. However, roughly half of cells in c1 also express exhaustion markers such as *Pdcd1* and *Ctla4* (16, 20).

Therefore, this population is likely a group of effector memory T cells with some transitioning into early exhaustion. C3 is comprised of CD8^+^ T cells that express genes associated with cell proliferation, such as *Mki67, Top2a*, and *Birc5* (21, 22) (Figure 3B, Supplementary Figures 3B). C3 also expresses high levels of *Runx3, Gzmb,* with a small percentage of cells expressing *Pdcd1* and *Ctla4*. *Runx3* is a key regulator of tissue-resident memory CD8+ T cell differentiation and homeostasis, while the other genes are associated with effector and early exhaustion CD8^+^ T cells (23). C5 is another cluster of CD8+ T cells that also express high levels of *Runx3, Cdca7, Stmn1, Txn1*, *Ifng,* and granzyme-related genes (Figure 3B, Supplementary Figures 3B). These genes are associated with higher immune cell infiltration, as well as proliferative CD8^+^ T cells (24–28). However, about 50% of the cells in c5 also express inhibitory receptors and transcription factors such as *Pdcd1* and *Ctla4*, which indicate that some of these cells are transitioning into an exhausted state (16). C7 expresses *Ifng, Gzmk, Gzmb*, and *Ly6c2*, suggesting that this cluster contains mostly effector memory T cells (29) (Figure 3B, Supplementary Figures 3B). Taken together, these data illustrate that there are increased effector/memory CD8^+^ T cells in the tumor after combination treatment with CpG and RT. In addition, these CD8^+^ T cells express relatively low levels of genes associated with exhaustion, usually less than 50% of T cells within a cluster express *Pdcd1* and *Ctla4*, which are also usually upregulated in activated T cells. Other common genes associated with exhaustion such as *Lag3, Havcr2,* and *Tox* are hardly expressed in these intratumoral T cells (Supplementary Figure 3B) (20). These T cells are also highly proliferative, indicating that CpG+RT attracts CD8^+^ T cells, which are not yet exhausted, into the tumors.

The population sizes of NK/ILC/NKT cells (c0 and c8 expressing *Ncr1*), Tregs (c4 expressing *Foxp3*), γδT cells (c9 expressing *Tcrg-c1*), T helper type 2 (Th2) cells (c10 expressing *Tbc1d4* and *Tnfsf8*), and a population of myeloid cells (c12 expressing *Cd74* and *H2-Aa*) did not differ significantly between the four treatment groups (Figure 3B, Supplementary Figures 3B, 3C) (28, 30–33). The three populations that were lower after CpG+RT compared to the control group are c2, c6, and c11 (Supplementary Figure 3C). C2 differentially expresses *Cd4, Lef1, S1pr1*, and *Ccr7*, which are genes generally associated with naive CD4^+^ T cells (34–38). C6 expresses *Cd8a, Lef1, Ccr7*, and *Il7r*, which are usually associated with naive CD8^+^ T cells (39). C11 is a cluster with very few cells that expresses *Slamf6, Eomes*, and *Foxp1*, which are often associated with exhausted or naïve T cells (Figure 3B, Supplementary Figures 3B). Taken together, these data indicate that CpG+RT resulted in increased infiltration of CD8^+^ T cells, with these cells displaying a range of transcriptional profiles indicative of effector memory, proliferation, and early exhaustion states. However, naive CD4^+^ and CD8^+^ T cells were more abundant in untreated tumors, suggesting a shift in the immune landscape with CpG+RT.

### Bulk RNA-seq demonstrated a similar increase in CD8^+^ T cells after CpG+RT treatment and specific T cell clonal expansion

To gain insight into the overall transcriptional differences in the tumor and immune microenvironments, we performed bulk tumor RNA-seq on an independent cohort of p53/MCA tumors harvested 6 days after RT and 3 days after CpG treatment or their respective controls. We then performed digital cytometry on the bulk tumor RNA-seq dataset using CIBERSORTx to estimate the abundance of 22 different immune cell populations in the tumor microenvironment (40). CIBERSORTx results from the bulk tumor RNA-seq dataset support the findings from CyTOF, CD8 immunohistochemistry, and single-cell RNAseq that CD8^+^ T cells increase significantly after treatment with CpG and RT (Figures 2C, 3D, 3E, 5B). These results from multiple orthogonal assays are consistent with the hypothesis that the superior treatment effect of combination treatment is mediated through influx, activation and proliferation of CD8^+^ T cells.

Given that we observed activation and proliferation of CD8^+^ T cells, we next evaluated whether there was clonal T cell expansion. T cell receptor (TCR) clonality analysis was conducted on the bulk RNA-seq data set and revealed that tumors treated with CpG+RT have the highest S-entropy score, indicating increased infiltration of different clones of T cells into the tumor (Figure 4A). This is consistent with our findings of increased CD8^+^ T cells in tumors after combination therapy (Figures 2 and 3). TCR clonality assessment also demonstrated that sarcomas treated with combination treatment had a lower evenness score when compared to tumors receiving control treatment or CpG alone, possibly due to preferential tumor-antigen specific T cell expansion after treatment rather than pan-T cell proliferation (Figure 4B).

**Fig 4.**
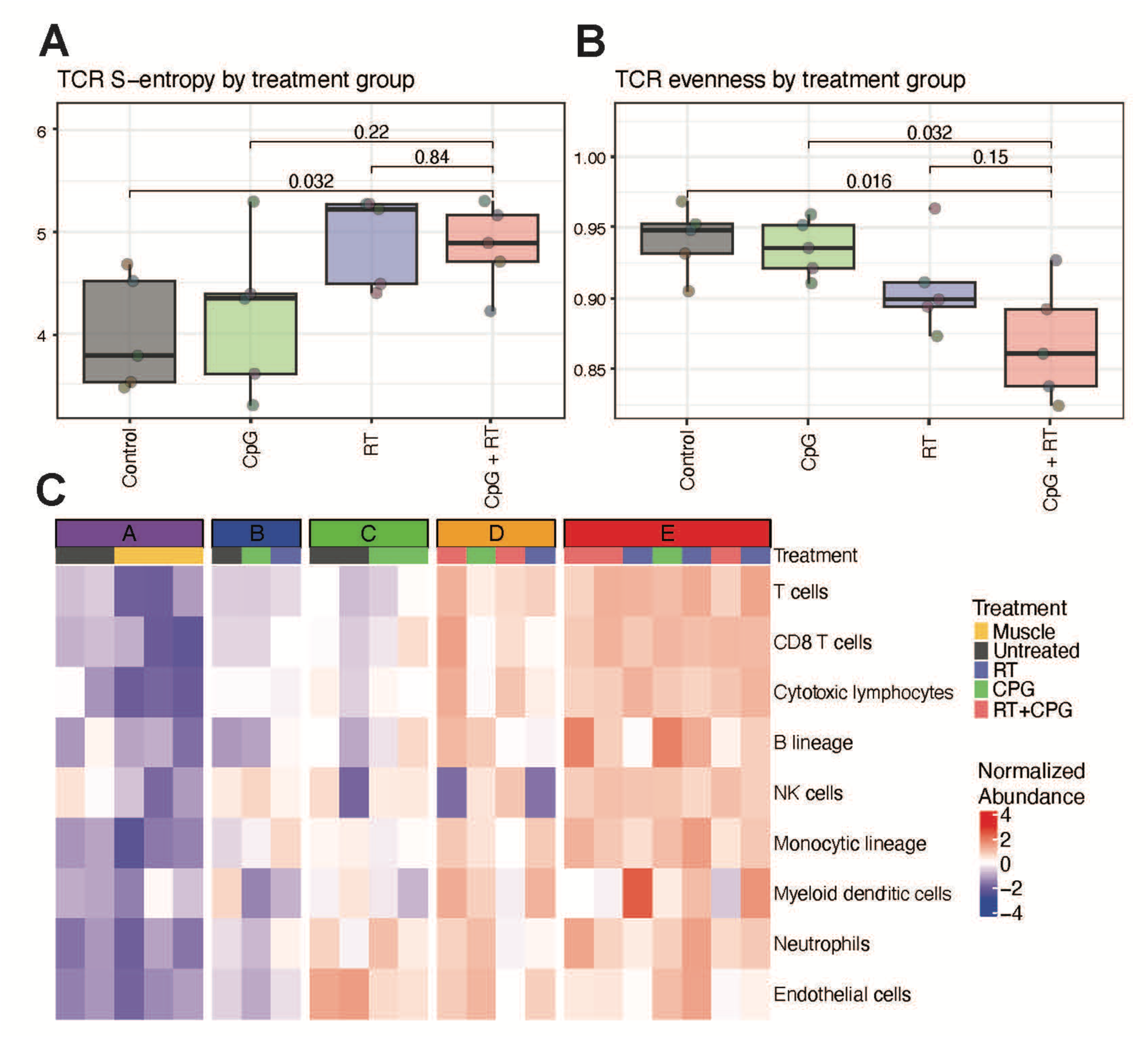
Treatment with RT + CpG promotes tumor-antigen specific clonal expansion of T cells and tumor immune microenvironment remodeling. **A.** p53/MCA sarcoma develops at the injection site about 7-11 weeks after induction. Mice were treated with CpG ODN or GpC dinulcotides control and 0 or 20 Gy when tumors reached >180 mm^3^. Sarcomas received control GpC dinucleotides with 0 Gy (*n* = 5), CpG ODN alone (*n =* 5), control GpC dinucleotides with 20 Gy (*n =* 5), or CpG ODN with 20 Gy (*n =* 5). Shannon entropy calculated from the abundance of TCR sequences captured by TCR sequencing and stratified by treatment group. Increasing entropy indicates reduced uniformity of TCR sequences. P values were calculated using two-sided Wilcoxon tests. **B.** TCR evenness is Shannon entropy normalized by species richness. **C**. Immune-based classification of murine primary sarcomas. Sample sizes: muscle control (n=3), tumor control (n=5), CpG ODN (n=5), RT (n=5), RT+CpG ODN (n=5).

Petitprez and colleagues described an immune-based classification (Classes A-E) of STS based on the composition of the tumor microenvironment (41). Sarcoma immune classes (SIC) D and E are associated with improved survival, and SIC E is associated with a higher response rate to anti-PD-1 treatment. Using the bulk RNA-sequencing, we assigned the murine p53/MCA tumors from each treatment group to an SIC based on the method described by Petitprez et al (Fig 4C). Mouse sarcomas treated with CpG and RT were all assigned to SIC D and E while sarcomas with no treatment were assigned to the less-inflamed SICs A, B, and C. Using CIBERSORTx and data from TCGA, we compared the immune infiltration in human UPS samples from each SIC with murine p53/MCA tumors from each treatment group (Fig 4C). Similar to SIC D and E human tumors, mouse sarcomas treated with CpG and RT had the highest immune infiltration, further demonstrating the similarity between these tumors. It is interesting to note that even though mouse sarcomas treated with combination therapy closely resembled human SIC D and E tumors, in our experimental system they did not respond to anti-PD1 treatment alone or in any combination with RT, CpG and/or OX-40 agonist antibody (Supplementary Figure 6A). We included OX-40 because it was previously reported to act with CpG to stimulate the immune response in an autochthonous mouse model of breast cancer (42), but in our model system OX40 was not active.

### Combination treatment of CpG and radiotherapy promotes myeloid cell remodeling and upregulates expression of MHC-I and MHC-II

We consistently observed an influx of CD8^+^ T cells in sarcomas treated with CpG and RT through CyTOF, scRNA-seq, and bulk tumor RNAseq data. However, TLR9 is the canonical receptor for CpG, and it is constitutively expressed by B cells and plasmacytoid dendritic cells rather than T cells. Therefore, we next used the CyTOF data to analyze TLR9-expressing antigen-presenting cells as they could regulate the profound CD8^+^ T cell proliferation and trafficking into the tumor after CpG+RT. We observed that TLR9 protein expression is upregulated in CpG+RT-treated tumors through CyTOF, especially in dendritic cells (DCs) and macrophages (Supplementary Figure 7A). We also observed significant upregulation of major histocompatibility complex (MHC) class I and class II proteins on antigen presenting cells after treatment with CpG and RT through CyTOF (Figure 5A, Supplementary Figure 7A). A similar increase in intratumoral CD11c+ DCs was also observed with CpG+RT compared to control (Figure 5A, Supplementary Figure 7A). These results support the notion that an increase in DCs and proteins associated with antigen presentation pathways promote the activation of CD8^+^ T cells after CpG+RT and thus enhanced tumor killing.

**Fig 5.**
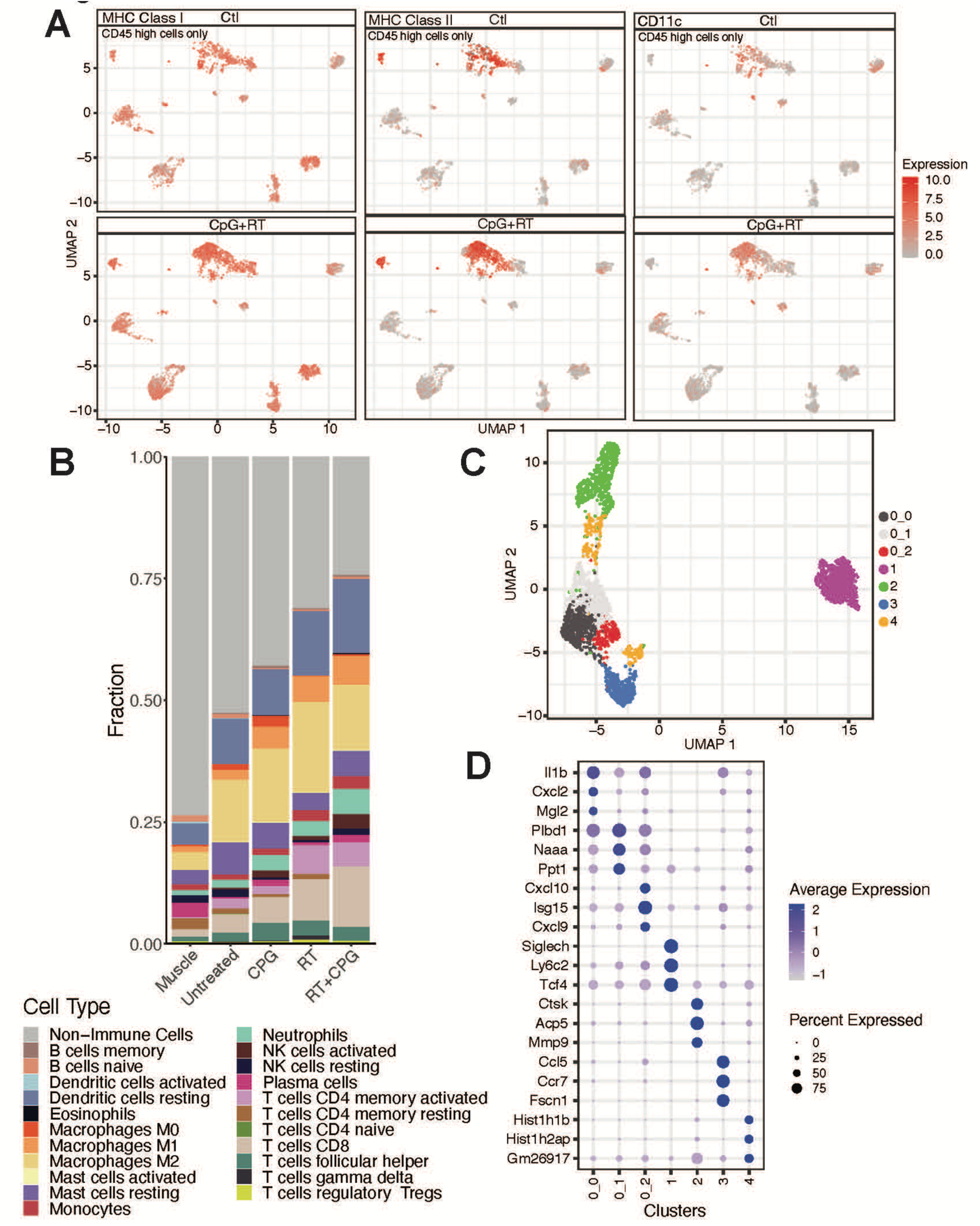
Treatment with RT + CpG promotes intratumoral myeloid cell remodeling. **A.** UMAP plot of CyTOF clustering for CD45hi cells from Ctl (GpC dinucleotides) and CpG+RT treatment groups. MHC-I, MHC-II, and CD11c expression are highlighted in red. **B.** Immune cell composition of all treatment groups from Bulk RNA-seq. **C.** UMAP plot of dendritic cells subclustering **D.** Bubble plot of top 3 differentially expressed genes in each of the DC subclusters. The shades of color are correlated with levels of expression. The sizes of circles are correlated with percentage of cells in that cluster that express the gene of interest. Sample sizes: tumor control (n=5), CpG ODN (n=5), RT (n=5), RT+CpG ODN (n=5).

CIBERSORTx results also demonstrate expansions of M1 macrophages and activated DCs after combination treatment with CpG and RT (Figure 5B).

Unsupervised clustering analysis of DCs resulted in the identification of 7 subpopulations (Figure 5C), which we compared to publications for cell-type identification based on differential gene expression (Figure 5D, Supplementary Figure 8, Supplementary Table 2). C0_0 is likely a group of proinflammatory conventional type 2 dendritic cells (cDC2s) that express high *Mhc-II, Il1b, Cd14,* and *Tnf*, as previously identified by Cheng and colleagues (43). C0_1 closely resembles conventional type 1 dendritic cells (cDC1s) with differential expression of *Itgae, Xcr1,* and *Clec9a* (43). CD103+ cDC1s have been shown to transport intact antigens to the tumor-draining lymph nodes and activate CD8^+^ T cells. C0_2 is characterized by increased expression of chemokine and interferon-inducible genes, such as *Cxcl10*, *Ifit1*, and *Isg15*, which were previously identified as cDC2_Isg15 (43).

C1 clustered farther apart from all other DCs and expresses signature genes representing plasmacytoid dendritic cells (pDC), such as *Siglec-h, Ly6c2, Bst2, Ptprcap,* and *Tcf4* (Figure 5D, Supplementary Figure 8) (44, 45). C3 resembles migratory DCs with high expression of *Ccl5, Ccr7, Ccl22,* and *Cacnb3* (46–48). None of these populations have significant changes in size based on treatment group (Supplementary Figure 7B).

C2 is likely a population of monocyte-derived DCs that express *Npr2* and *Ctsk*, which play important roles in DC maturation, enhanced DC-T cell interactions, and promoting TLR9-induced cytokine production (49–51). It is worth noting that C2 almost completely disappears after treatment with RT or CpG+RT (Supplementary Figure 7B). Since both bulk tumor RNA-seq and CyTOF data demonstrated increased infiltration of DCs after combination treatment, it is plausible to hypothesize that C2 migrated to the lymph nodes for antigen presentation after CpG+RT. However, it is also possible that C2 was sensitive to radiation and was eliminated after treatment. C4 is another DC population that decreased after combination treatment (Figure 5D). C4 expresses *Itgam* (CD11b), but not *Itgae* (CD103), which is often associated with migratory cDCs that travel to lymph nodes for antigen presentation (52, 53). These findings suggest that CpG+RT treatment promoted DC maturation by upregulating genes that are associated with antigen presentation, such as MHC-I and MHC-II. The combination treatment also induced upregulation of genes associated with lymph node trafficking and interactions with T cells.

### Lymphocytes, especially CD8^+^ T cells, are crucial in mediating the anti-tumor effects of CpG and radiotherapy in vivo

To evaluate whether the CpG+RT treatment effect is indeed mediated through the adaptive immune system *in vivo*, we induced p53/MCA tumors in *Rag2^-/-^;γc^-^* (male); *Rag2^-/-^;γc^-^*^/-^ (female) and their littermate controls *Rag2^+/-^;γc^+^* (male); *Rag2^+/-^;γc^+/-^* (female) (Figure 6A). Since the *γc* gene is X-linked, the genotypes for littermate controls are different between males and females. *Rag2^-/-^;γc^-^* (male) and *Rag2^-/-^;γc^-^*^/-^ (female) mice are incapable of generating functional B cells, T cells or NK cells. When tumor volume reached 70-150 mm^3^, mice were randomized to receive 0 Gy or 20 Gy RT (Day 0) and CpG or control (GpC dinucleotides with the positions of cytosine and guanine reversed relative to the phosphate linker) (Day 3 and Day 10) (Figure 6B). The increase in time to tumor quintupling with CpG+RT compared to control was similar for heterozygous littermate controls that retained functional B cells, T cells or NK cells (Figures 6C and D) and 129/SvJae mice (Figure 1C). However, this treatment effect was lost in homozygous *Rag2γc* double knockout (DKO) mice (*Rag2^-/-^;γc^-^* (male); *Rag2^-/-^;γc^-^*^/-^ (female)) (Figures 6E and F), suggesting that the adaptive immune system plays a crucial role in facilitating the anti-tumor effects of CpG and RT.

**Fig 6.**
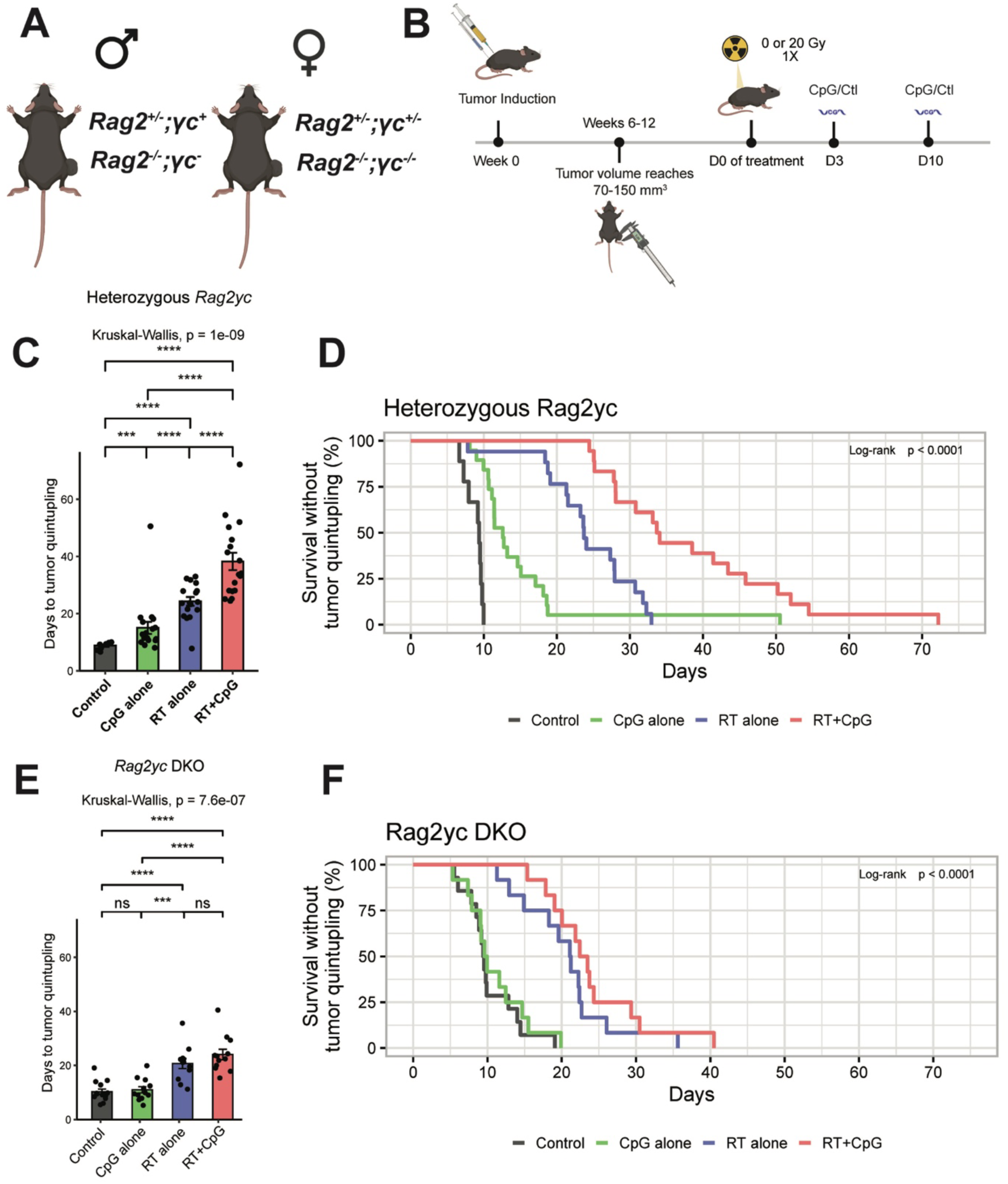
Lymphocytes mediate the anti-tumor effects of the combination treatment RT + CpG. A. Primary sarcoma initiation by intramuscular injection of Adeno-Cas9-sgp53 and MCA. **B**. Autochthonous sarcoma develops at the injection site about 7-11 weeks after induction. Mice were treated with CpG ODN or control GpC dinucleotides and 0 or 20 Gy when tumors reached >70 mm^3^. **C.** Heterozygous Mice (Rag2+/-;yc+ or Rag2+/-;yc+/-) with p53/MCA sarcomas received control GpC dinucleotides with 0 Gy (black, *n* = 9), CpG ODN alone (green, *n =* 17), control GpC dinucleotides with 20 Gy (blue, *n =* 17), or CpG ODN with 20 Gy (red, *n =* 18). Figure shows time to tumor quintupling (days). **D**. Heterozygous mice (Rag2+/-;yc+ or Rag2+/-;yc+/-) with p53/MCA sarcomas received control GpC dinucleotides with 0 Gy (black, *n* = 9), CpG ODN alone (green, *n =* 17), control GpC dinucleotides with 20 Gy (blue, *n =* 17), or CpG ODN with 20 Gy (red, *n =* 18). Figure shows time to tumor quintupling (days). **E**. Homozygous mice (Rag2-/-;yc- or Rag2-/-;yc-/-) with p53/MCA sarcomas received GpC dinucleotides control with 0 Gy (black, *n* = 14), CpG ODN alone (green, *n =* 12), GpC dinucleotides control with 20 Gy (blue, *n = 12*), or CpG ODN with 20 Gy (red, *n =* 12). Figure shows time to tumor quintupling (days). **F**. Homozygous mice (Rag2-/-;yc- or Rag2-/-;yc-/-) with p53/MCA sarcomas received GpC dinucleotides control with 0 Gy (black, *n* = 14), CpG ODN alone (green, *n = 12),* GpC dinucleotides control with 20 Gy (blue, *n = 12*), or CpG ODN with 20 Gy (red, *n =* 12). Figure shows time to tumor quintupling (days). Kruskal-Wallis Test was used for the group comparison, while the Wilcoxon test was selected for the pair-wise comparisons. * shows the significance of the P Value (***: p <= 0.001, ****: p <= 0.0001).

We next directly tested if CD8^+^ T cells are necessary for the treatment effects observed with CpG+RT through CD8^+^ T cell depletion in 129/SvJae mice with p53/MCA sarcomas. When tumors reached 70-150 mm^3^, mice received intraperitoneal injections of isotype control or anti-CD8 antibodies (Abs) on the same day as RT or sham RT (Figure 7A). Isotype control or CD8-depleting Abs are repeated every 3-4 days until euthanasia upon reaching the humane endpoint (Figure 7B). Tumor growth delay was observed with CpG+RT in the isotype control group (Figures 7B and C), but the growth delay with combination treatment was not observed in the CD8-depleted mice (Figures 7D and 7F). These results demonstrate the essential role of CD8^+^ T cells in tumor growth delay induced by CpG and RT.

**Fig 7.**
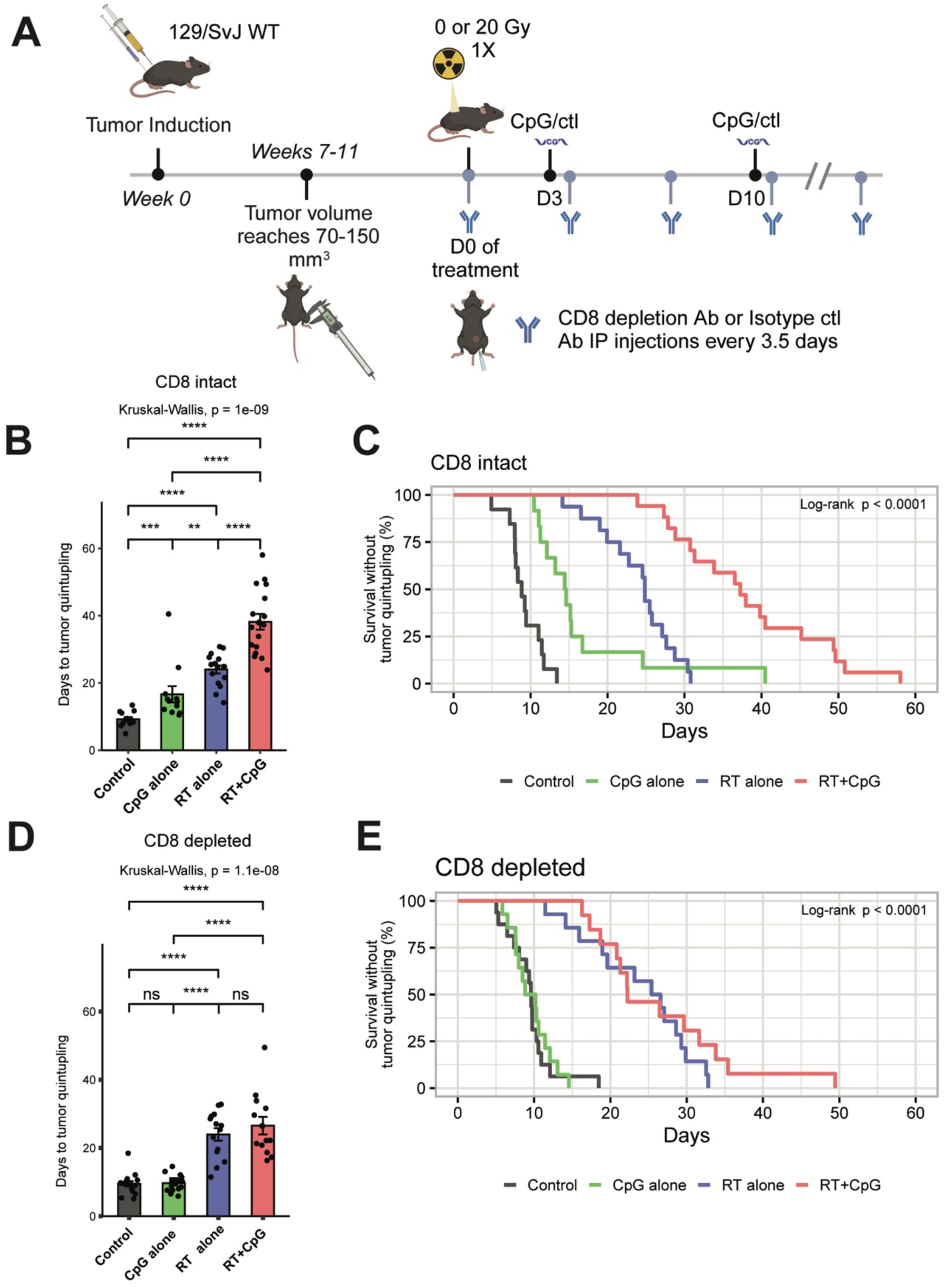
CD8 T cells are required for the treatment effects of RT + CpG. A. Primary sarcoma initiation by intramuscular injection of Adeno-Cas9-sgp53 and MCA. Autochthonous sarcoma develops at the injection site about 7-11 weeks after induction. Mice were treated with CpG ODN or control GpC dinucleotides and 0 or 20 Gy when tumors reached >70 mm^3^. Mice received intraperitoneal CD8 isotype control or CD8 depletion antibodody (Ab) on the same day tumors received RT. CD8 isotype control or CD8 depletion Ab are repeated every 3.5 days until tumor size reached humane end point. **B**. 129/SvJ mice with p53/MCA sarcomas, injected with CD8 isotype control Ab, received GpC dinucleotides control with 0 Gy (black, *n* = 13), CpG ODN alone (green, *n =* 12), GpC dinucleotides control with 20 Gy (blue, *n =* 15), or CpG ODN with 20 Gy (red, *n =* 14). **C**. 129/SvJ mice with p53/MCA sarcomas, injected with CD8 isotype control Ab, received GpC dinucleotides control with 0 Gy (black, *n* = 13), CpG ODN alone (green, *n =* 12), GpC dinucleotides control with 20 Gy (blue, *n =* 15), or CpG ODN with 20 Gy (red, *n =* 14). **D**. 129/SvJ mice with p53/MCA sarcomas, injected with CD8 depleting Ab, received GpC dinucleotides control with 0 Gy (black, *n* = 16), CpG ODN alone (green, *n =* 14), GpC dinucleotides control with 20 Gy (blue, *n =* 13), or CpG ODN with 20 Gy (red, *n =* 11). **F**. 129/SvJ mice with p53/MCA sarcomas, injected with CD8 depletion Ab, received GpC dinucleotides control with 0 Gy (black, *n* = 16), CpG ODN alone (green, *n =* 14), GpC dinucleotides control with 20 Gy (blue, *n =* 13), or CpG ODN with 20 Gy (red, *n =* 11). Figure shows time to tumor quintupling (days). Kruskal-Wallis Test was used for the group comparison, while the Wilcoxon test was selected for the pair-wise comparisons. * shows the significance of the P Value (***: p <= 0.001, ****: p <= 0.0001).

## Discussion

Many studies have explored synergistic effects between immunotherapies and radiation therapy in sarcoma (3, 6, 7). However, most preclinical experiments are conducted with xenograft or transplanted models in which the tumor does not coevolve with the host immune system. A number of studies have shown that autochthonous cancer models, which develop under immunosurveillance, more faithfully recapitulate the microenvironment of human cancers (54, 55). Therapeutic approaches that elicit impressive survival benefits in transplanted tumor models may fail when translates into clinical trials (10, 56, 57). In this study, we addressed the limitations of transplant tumor models by utilizing the high mutational load autochthonous p53/MCA murine sarcoma model, which allows the tumor to develop under surveillance of an intact immune system (8, 9). Our results demonstrate an enhanced radiation response of primary sarcomas treated with intratumoral CpG as measured by tumor growth delay, which is mediated through the activation and expansion of intratumoral CD8^+^ T cells. Our work suggests that the combination of radiation therapy with a TLR9 agonist, such as CpG, warrants evaluation in a clinical trial of sarcomas and perhaps other cancers.

Mechanistically, our work indicates that combination treatment with CpG and RT enhances the activation and proliferation of intratumoral CD8^+^ T cells, as demonstrated by CyTOF, IHC, scRNAseq, and bulk tumor RNAseq. These CD8^+^ T cells express high levels of granzymes and IFNγ, indicating that they are activated and capable of cellular cytotoxicity. Furthermore, the majority of these T cells are also in the S or G2-M phases of the cell cycle, indicating active proliferation. The TCR clonality analysis further supports this model, revealing not just an increase in CD8^+^ T cell numbers, but also specific infiltration of certain T cell clones, which suggests targeted tumor-antigen-specific T cell response rather than a general proliferation of T cells. Additionally, our findings show that p53/MCA sarcomas in mice treated with CpG and RT exhibit immune profiles similar to SIC classes D and E, which are associated with better survival outcomes and response rates to anti-PD1 therapy in patients with STS. Furthermore, CD8^+^ T cell depletion in the murine model abrogated the treatment effect of CpG+RT, which establishes a critical role of CD8^+^ T cells in mediating the treatment effects of this combination therapy.

Future functional studies are needed to determine the underlying mechanism responsible for CD8^+^ T cell activation and trafficking to the tumor after CpG and RT. Sagiv-Barfi and colleagues explored the combined therapeutic effects of the administration of local TLR9 agonist with systemic anti-OX40 agonist in murine models with spontaneous mammary gland tumors. They observed significant tumor burden reduction not only at the TLR9 agonist injection site, but also at distant tumor sites (42). They reported upregulated OX40 expression on CD4^+^ T cells after CpG treatment and superior treatment response when anti-OX40 was added to intratumoral CpG injections (42). However, we did not observe increased OX40 expression after CpG+RT (Supplementary Figure 6B). Nor did we observe a synergistic treatment effect with anti-OX40 in the p53/MCA sarcoma model (Supplementary Figure 6A).

There were some differences in the cell population changes after CpG+RT treatment in scRNA-seq data versus bulk tumor RNA-seq, which might stem from different rates of mRNA recovery in the sample preparation process. For example, both bulk tumor RNA-seq and CyTOF data demonstrated an increase in DC populations after combination treatment with CpG+RT. However, scRNA-seq generally showed a decrease in the number of infiltrating DCs after combination treatment suggesting a technical limitation for detecting DCs with scRNA-seq in our experiments. 10X Genomics reported about 30-32% of mRNA transcripts are captured per cell utilizing the Single Cell 3’ reagent chemistry v3 (58). However, Qiagen reports more than 90% mRNA recovery from tissues utilizing their mRNA extraction kit (59). Therefore, differences in mRNA recovery could partly account for discrepancies in dendritic cell populations observed between bulk tumor RNA-seq and scRNA-seq data.

Overall, CpG appears to be an excellent candidate for treatment of patients with STS due to its ease of local administration and favorable safety profile (60, 61). No apparent toxicity was observed in mice treated with CpG+RT. Furthermore, our *in* vivo studies with p53/MCA sarcomas demonstrated that using TLR9 agonists in conjunction with radiotherapy significantly outperformed the individual treatments or no treatment in terms of delaying tumor progression. Seo and colleagues recently published results from a clinical trial utilizing intratumoral injection of a TLR4 agonist and radiotherapy to treat 12 patients with metastatic sarcoma (62). All tumors treated with the combination therapy achieved durable local control, while tumors receiving RT alone or no treatment showed limited or no response. Similarly, our *in vivo* studies in mice with p53/MCA sarcomas demonstrated that intratumoral injection of CpG as a TLR9 agonist in conjunction with radiotherapy significantly improved response compared to either treatment alone as measures by tumor growth delay. Our results with CpG+RT and the initial clinical trial of a TLR4 agonist with RT demonstrate the potential effectiveness of TLR agonists and radiotherapy in treating sarcomas. Furthermore, unlike other immunotherapies, such as immune checkpoint inhibitors that aim to reverse the exhaustion state of T cells, TLR9 agonist combined with RT causes activated CD8^+^ T cells that are not yet exhausted to infiltrate the tumor and enhance the radiation response.

In summary, we find that intratumoral CpG enhances the response of primary sarcomas to radiotherapy by increasing CD8+ T cell infiltration. This study supports translating the therapeutic approach of radiotherapy with CpG or another TLR9 agonist into clinical trials for patients with soft-tissue sarcomas.

## Methods

### Experimental models details

All animal studies were performed in accordance with protocols approved by the Duke University Institutional Animal Care and Use Committee (IACUC) and adhered to the NIH Guide for the Care and Use of Laboratory Animals. The *Rag2^-/-^ γc^-/-^mice* and their littermates were purchased from Jackson Laboratory and bred at Duke University with a mixed BALB/cAnNTac and 129S4/SvJae background. Wild type 129S4/SvJae mice used in this study were also purchased from Jackson Laboratory and bred at Duke University. To minimize the effects of sex and genetic background, male and female mice and age-matched littermate controls were used for every experiment so that potential genetic modifiers would be randomly distributed between experimental and control groups.

### Sarcoma Induction and Treatment

Primary p53/MCA sarcomas were generated in 129S4/SvJae and *Rag2^-/-^ γc^-/-^* or littermate control mice between 6 and 12 weeks old by intramuscular injection of adenovirus expressing Cas9 and sgRNA targeting *Trp53* (Adeno-p53-sgRNA; Viraquest) into mice as previously described (10). Twenty-five microliters of adenovirus were mixed with 600 µL DMEM (Gibco) and 3 µL 2 M CaCl2, then incubated for 15 min at room temperature prior to injection. Fifty microliters of the prepared mixture were injected into the gastrocnemius muscle, followed by injection of 300 µg MCA (Sigma) resuspended in sesame oil (Sigma) at 6 µg/µL.

For tumor growth delay studies in 129S4/SvJae and *Rag2^-/-^ γc^-/-^* or littermate control mice, mice were randomized to treatment groups when tumors reached 70–150 mm^3^ (Day 0, D0). Tumors were monitored three times weekly by caliper measurements in two dimensions until one dimension of the tumor reached 15mm. Mice received one dose of 0 or 20 Gy of image-guided radiation therapy to the tumor-bearing hind limb on D0. Mice were anesthetized with 2% isoflurane and 98% oxygen at 2 L/min and held on the specimen positioning stage of a Micro-CT on a Small Animal Radiation Research Platform (First half of tumor quintupling studies and scRNAsesq and bulk tumor RNAseq experiments were conducted utilizing SmART+, Precision Inc. Second half of tumor quintupling studies and CyTOF experiments were conducted using SARRP, Xstrahl Inc.). The right hind limb was identified using micro-CT guided fluoroscopy (60 kVp, 0.8 mA X-rays using a 1 mm Al filter). Irradiations were performed using parallel-opposed anterior and posterior X-rays were delivered via 20 mm x 20mm collimators (220 kVp, 13 mA X-rays using a 0.15 mm Cu filter) with a dose of 20 Gy of radiation prescribed to mid-plane delivered in a single, unfractionated dose.

50ul of CpG or GpC Control (Ctl) was injected intra- and peritumorally on D3 and D10. CpG (InvivoGen, ODN 1826) or GpC Ctl with the positions of cytosine and guanine reversed relative to the phosphate linker (InvivoGen, ODN 2138) was diluted in endotoxin-free water at 1mg/ml. Antibodies were administered starting on D0 (on the same day as RT) by intraperitoneal injection of 200 µL per dose at 1mg/mL diluted in PBS. Anti-CD8 (BioXCell, BE0061) or isotype control (BioXCell, BE0090) antibodies were injected every 3-4 days for the duration of the experiment. Anti-PD-1 (BioXCell, BE0146) or isotype control (BioXCell, BE0089) were injected on D3, D7, D10. Anti-OX40 (Bristol Myers Squibb) or isotype control (Bristol Myers Squibb) were injected on D3 and D10.

### Mass cytometry

#### Tumor harvest and dissociation

Tumors were dissected from mice, minced, and digested using the Miltenyi Biotec tumor dissociation kit (mouse, tough tumor dissociation protocol) for 40 min at 37°C. Cells were then strained through a 70 µm filter and washed with Maxpar Cell Staining Buffer (CSB) (Standard Bio Tools). Red blood cells (RBCs) were lysed using ACK lysis buffer (Lonza). Cells were then washed and resuspended in Maxpar PBS for cell counting using Trypan Blue (Thermo Fisher).

#### CyTOF Staining

For custom-conjugated antibodies, 100 µg of antibody was coupled to Maxpar X8 metal-labeled polymer according to the manufacturer’s protocol (Standard Bio Tools). After conjugation, the metal-labeled antibodies were diluted in Antibody Stabilizer PBS (Candor Bioscience) for long-term storage according to the manufacturer’s protocol. After tumor dissociation and RBC lysis as described above, three million cells per sample were transferred to 5 mL round-bottom tubes (Corning). Cells were incubated with 300 µL of Cell-ID Cisplatin-195Pt (Standard Bio Tools) diluted 1:8000 in Maxpar PBS (Standard Bio Tools) for 5 min at room temperature, then washed with CSB (Standard Bio Tools). Samples were incubated with 1.5 µg TruStain FcX PLUS Blocking Reagent (Biolegend) for 10 min at room temperature, then 56.55 µL extracellular antibody cocktail was added and incubated for 30 min at room temperature. The final staining volume is 130 µL including residual CSB from the wash, FcX PLUS blocking reagent, CSB, and antibody cocktail. Cells were washed twice with CSB, then fixed and permeabilized with Foxp3/Transcription Factor Fixation/Permeabilization Buffer (eBioscience) overnight at 4°C. The next morning, cells were washed twice with permeabilization buffer (eBioscience). Fifty microliters of intracellular antibody cocktail in permeabilization buffer was added and incubated for 30 min at room temperature, followed by two washes with permeabilization buffer. Cells were fixed in 1.6% methanol-free paraformaldehyde (PFA) (Thermo Fisher) diluted with Maxpar PBS (Standard Bio Tools) for 10 minutes at room temperature. Samples were incubated for 1 hour in Maxpar Fix and Perm Buffer (Standard Bio Tools) with 1 mL of 157.2 nM Cell-ID Intercalator (Standard Bio Tools) containing 191Ir and 193Ir. After staining, samples were centrifuged and resuspended in 100 µL of residual Intercalator/Fix and Perm Buffer. Cells were transferred to 1.5 mL microcentrifuge tubes and stored at −80°C. On the day of acquisition, cells were washed once with CSB, once with Cell Acquisition Solution (CAS) (Standard Bio Tools), then filtered and diluted in CAS containing 10% EQ Calibration Beads (Standard Bio Tools) at 0.5 million cells per mL before acquisition on a mass cytometer (Helios).

#### CyTOF data analysis

Mass cytometry data were analyzed using Standard Bio Tools CyTOF software (v7.0). Individual samples were gated in Cytobank to exclude beads, debris, dead cells, and doublets for further analysis. For each experimental group (treatment), cells from 8 to 11 tumors per group were manually gated to identify specific populations.

#### CyTOF dimension reduction and clustering

Expression data from all .fcs files were log2 transformed. We used unbiased clustering with FastPG v0.0.8 to identify clusters of cells in the expression data (63). A k of 50 was used and all cells were included (CD45 high & low) in the clustering step. Dimension reduction was performed on the log2 expression data using both PCA and UMAP approaches. The statistical significance of markers across treatment groups was determined using a Wilcoxon rank-sum test.

### Bulk tumor RNA sequencing of p53/MCA sarcomas in 129S4/SvJae mice

#### RNA extractions

Tumor specimens and matched muscle control were harvested and stored in RNALater (Ambion) at −80°C until all samples were collected. RNA extractions from each sample were performed using RNeasy Fibrous Tissue Mini Kit (Qiagen). Extracted total RNA quality and concentration were assessed on a NanoDrop Spectrophomoter (Thermo Fisher Scientific). RNA-seq libraries were prepared using TruSeq Small RNA Library Preparation Kits (Illumina) following the manufacturer’s protocol. RNA sequencing was performed on an Illumina Novoseq in 151 base pair, paired-end configuration. Greater than 50,000 reads per cell were collected per the manufacturer’s recommendation.

#### Sequence alignment and sarcoma immune class labels

Mouse FASTQ files were aligned to the GRCm38 reference genome using Salmon (64). Mouse genes were mapped to their corresponding human orthologs using the ‘biomaRt’ R package (65). The expression matrix was restricted to protein-coding genes and expression levels per gene were summarized as transcripts per million (TPM). Pre-processed bulk RNA-seq profiles from human UPS from TCGA were downloaded and scaled to TPM. Sarcoma immune class labels were obtained from the authors (41, 66). *CIBERSORTx RNA sequencing analysis*

Cell type proportions were inferred using CIBERSORTx with two previously published signature matrices (LM22 and TR4) (40). LM22 encompasses 22 distinct human immune cell subsets, and TR4 comprises epithelial, fibroblast, endothelial, and immune cells (40, 67). Bar plots and heatmaps were visualized with the ‘ggplot2’ and ‘ComplexHeatmap’ R packages, respectively (68). For the heatmaps, abundances were normalized to mean zero and unit variance. P values were calculated using two-sided Wilcoxon tests.

#### Assignment of mouse sarcomas to sarcoma immune classes

Sarcoma immune classes were assigned to normal muscle and mouse sarcomas in each treatment group as described previously (41). Briefly, abundance scores for T cells, CD8 T cells, cytotoxic lymphocytes, B cell lineage, natural killer cells, monocytic lineage, myeloid dendritic cells, neutrophils, and endothelial cells were determined from the bulk RNA sequencing using MCP-counter (69) and normalized across samples. Because the transcriptional profiles of the mouse tumors in this study are most similar to human undifferentiated pleomorphic sarcomas, we constructed centroids using the MCP-counter Z-scores for undifferentiated pleomorphic sarcomas from TCGA (66) based on the sarcoma immune class labels obtained from the authors (41). Mouse samples were assigned to the closest sarcoma immune class by evaluating the Euclidean distance to each centroid.

#### Gene differential expression analysis

Differential expression analysis was performed using the ‘DESeq2’ R package with treatment condition as the design variable (70). Expression of individual genes was visualized by plotting DESeq2 normalized counts. Volcano plots were generated using the ‘EnhancedVolcano’ R package. Pre-ranked gene set enrichment analysis for the hallmark gene sets was performed with the ‘fgsea’ R package using the negative log10 of the p-value multiplied by the direction of the log fold change for each gene (71, 72). Data were analyzed using R version 4.1.2.

#### Gene expression analysis for TCR clonality analysis

All FASTQ files were processed using a combination of STAR v2.7.10a and Salmon v1.2.0 with the mouse build GRCh38 - mm10 as a reference (64, 73). FASTQ files from the same sample, but different lanes, were merged with STAR’s fastq_channel_merged_paired function. Reads were mapped using STAR’s fastq_channel_merged_paired_star function. Picard and QC, count matrix assembly, and bam files were generated for each sample using both Picard v2.27.4, STAR v2.7.10a, and Salmon v1.2.0 (74).

#### TCR reconstruction and CDR3 metrics

TRUST4 v1.0.12 (Tcr Repertoire Utilities for Solid Tissue) were used to reconstruct TCR and BCR sequences from BAM files generated from the STAR/Salmon outputs (75). All parameters for TRUST4 were set to default. The final output from TRUST4 was summary matrices for each sample that included hypervariable complementarity-determining region 3 (CDR3) nucleotide and amino acid sequences, counts, frequencies, and V, D, and J chain names. Using the outputs from TRUST4, we calculated the following diversity metrics for all CDR3 species per treatment group: Shannon entropy and evenness by sequence were used to compare TCR species across treatment groups (76, 77). Shannon entropy measures the richness and abundance of each species. Species evenness measures the uniformity of the species. The calculations for each metric given a single TCR species are as follows: Count proportion: species count÷total species count Species richness: nucleotide length Shannon entropy: -Σ(count proportion× log(count proportion)) Evenness by sequence: Shannon entropy÷ log(species richness)

### Single-cell RNA sequencing of p53/MCA sarcomas in 129S4/SvJae mice

#### Tumor harvest and dissociation

Tumors were dissected from mice, minced, and digested using the Miltenyi Biotec tumor dissociation kit (mouse, tough tumor dissociation protocol) for 40 min at 37°C. Cells were then strained through a 70 µm filter and washed with FACS buffer (HBSS (Gibco) with 5 mM EDTA (Sigma-Aldrich) and 2.5% fetal bovine serum (Gibco). RBCs were lysed using ACK lysis buffer (Lonza) and washed again with FACS buffer.

#### Fluorescence-activated cell sorting

Dissociated cells were prepared for fluorescence-activated cell sorting of CD45+ cells for single-cell RNA sequencing. Single-cell suspensions of tumor tissues were blocked with 1.5 µg TruStain FcX PLUS Blocking Reagent (Biolegend) for 10 min at room temperature then stained with Live/Dead dye (Zombie Aqua, Biolegend) and anti-mouse CD45 (APC-Cy7, Biolegend) for 25 min on ice. Live CD45+ cells were isolated for scRNA-seq using an Astrios (Beckman Coulter) sorter and resuspended in PBS with 0.04% BSA at a concentration of 1000 cells/µL for single-cell RNA sequencing.

#### Library preparation and sequencing

Single-cell suspensions from sorted live CD45+ cells were loaded on a Chromium Controller (10x Genomics) to generate single-cell beads in emulsion, and scRNA-seq libraries were prepared using the Chromium Single Cell 3′ Reagent Kits (v3.1 Single Index Kit), Chromium Next GEM Single Cell 3′ GEM, Library & Gel Bead Kit v3.1, Chromium Next GEM Chip G Single Cell Kit, and Single Index Kit T Set A (10x Genomics) following the manufacturer’s protocol. Single-cell barcoded cDNA libraries were qualified and quantified using Agilent Bioanalyzer. cDNA libraries were sequenced on an Illumina NextSeq 500 (Illumina). Read lengths were 26 bp for read 1, 8 bp for i7 index, and 98 bp for read 2. Ten thousand cells from each sample were sequenced with greater than 50,000 reads per cell as recommended by the manufacturer.

#### Analysis of scRNA-seq data

Raw FASTQ files were mapped to *Mus musculus* reference mm10 GRCm38 using CellRanger v6.1.1 (10X Genomics) with default parameters. Expression matrix assembly and calculation of cell metrics (gene counts, molecule counts, percent mitochondrial genes) were performed using Seurat v.4.1.0 (78). We removed low-quality cells that had total non-zero gene counts ≤500 and ≥5,000, total non-zero molecule counts ≤2,000 and ≥40,000, ≥10% mitochondrial gene presence, a hybrid doublet score ≥1.0 per SCDS default parameters (79), and had an ambient RNA contamination score ≥0.50 per decontamX default parameters (80). Count data was normalized using LogNormalize in Seurat with a scaling factor of 10,000. The top 10,000 variable features were then used to scale the data using Seurat’s ScaleData function. Although cell cycle genes were scored using CellCycleScoring in Seurat, no variables were regressed out upon scaling. Cell types were annotated using SingleR v.1.8.1 and the mouse immunological genome project dataset (Immgen) available in the celldex v.1.4.0 library as a reference (81). Dimension reduction was performed using Harmony v0.1.0 with RunHarmony set to theta of 2 and all other parameters set to default (82). Harmony reductions were used for all downstream UMAP and clustering calculations. To select a clustering resolution, we calculated the adjusted rand index (ARI) and selected the clustering resolution with the highest ARI. Additional sub-clustering was performed downstream on cell types individually, and T cell subtypes were annotated using the T cell atlas from projecTILs with filter.cells set to “TRUE” (83).

### Immunohistochemistry Staining

Parts of the sarcoma were preserved for IHC staining when tumors were harvested for bulk tumor RNA-seq. Small chunks of sarcoma were fixed in 10% formaldehyde overnight and then preserved in 70% ethanol until paraffin embedding. Formalin-fixed paraffin-embedded tissues were sectioned and fixed onto slides for staining. Tissue sections were then deparaffinated by xylene and rehydrated with a series of graded ethanol and tap water. Slides were cooked in a rice cooker with Antigen Unmasking Solution, Citric Acid Based (Vector Laboratories) for epitope retrieval. After washes with tap water and PBS, tumor sections were then incubated in normal goat serum for 30 minutes at RT and then with CD8 antibody (Cell Signal, 98941) at 4°C overnight. The following day, slides were washed with PBS and then incubated with biotinylated anti-Rabbit antibody (Vector Laboratories) for 30 minutes at RT. VECTASTAIN ELITE ABC-HRP Reagent (Vector Laboratories) was then added to tumor sections for another 30 minutes at RT. Mayer’s hematoxylin (Sigma-Aldrich) was applied to the slides for counter-staining. Tumor slides were dehydrated and sealed with coverslips. Ten fields per tumor slide were randomly selected and counted by an observer blinded to treatment at 40X magnification for the number of CD8^+^ T cells.

### Cytotoxicity study for CpG effect on tumor cells

#### Generation of sarcoma cell lines

We generated individual cell lines from three murine autochthonous p53/MCA sarcomas as described above. Tumors were dissected from the limb and dissociated by shaking for 45 min at 37°C in an enzyme mix from the tumor dissociation kit (Miltenyi Biotec, 130096730). Cell suspension was then strained through a 40 µm filter, washed in PBS, and plated for culture. Cell lines were maintained *in vitro* for 5 passages before coincubation with CpG in 96-well plates.

#### In vitro tumor proliferation assay with CpG through IncuCyte

1×10^4^ p53/MCA sarcoma cells were resuspended in 100µL of complete cell culture media. After cells adhere to the bottoms of the plate overnight, 100µl of 0.01µg/µl, 0.1µg/µl, and 1µg/µl of CpG and 0.9ul of caspase-3/7 dye (Sartorius, 4440) were added to the well for 72hrs incubation in the SpectraMax M5 and Lmax microplate reader. Images of the sarcoma cells acquired by the microplate reader were analyzed by the Incucyte^®^ Software and plotted using GraphPad Prism.

### Statistics and study design

Experiments were designed such that littermate controls were used for all experiments. For bar graphs, all data are presented as mean ± SEM. For comparison of time to tumor quintupling in tumor growth experiments, we performed a Kruskal-Wallis test (after all groups failed the Kolmogorow-Smirnov test for normality) on the time to tumor quintupling (days) for each sample stratified by treatment group. To identify differences between groups, we performed a pairwise Wilcoxon test. The * shows the significance of the P-value (*: p <= 0.05, **: p <= 0.01, ***: p <= 0.001, ****: p <= 0.0001, ns: p > 0.05). This was complemented by a Kaplan-Meier curve and a log-rank test to compare survival without tumor quintupling across treatment groups.

## Data availability

All sequencing data generated for this manuscript have been deposited in publicly accessible databases. The p53/MCA bulk tumor RNA-seq data generated in this study are available in the NCBI Gene Expression Omnibus (GEO) database under the accession code GSE252213. The p53/MCA single cell RNA-seq data generated in this study are available in the NCBI Gene Expression Omnibus (GEO) database under the accession code GSE252143. The mass cytometry data generated in this study are available in the flowrepository.org database under the ID code FR-FCM-Z74J. Code for scRNA-seq analysis in the manuscript can be found at https://github.com/Quant-Bio/kirsch_rad_cpg_immunotherapy. The remaining data are available within the Article, Supplementary Information or available from the authors upon request. Source data are provided with the paper. Correspondence and requests for materials should be addressed to D.G.K.

## Author Contributions

CS and DGK conceptualized the project. CS, CLK, AJW, EJW, YMM, and DGK developed the methodology. CS, CLK, AS, VMP, RDH, KER, JLM, SRR and EJM conducted statistical analysis. CS, CLK, MP, LL, NTW, YM, WF, BP, ALL, and JEH performed the investigation. CS and DGK wrote the original draft of the manuscript. CS, AS, VMP, MS, AJW, EJM, YMM, and DGK reviewed and edited the manuscript. CS was responsible for visualization. DGK supervised the project. CS and CLK are co-first authors. CS is listed first due to her writing contributions.

## Acknowledgements

We thank Malay Haldar for advice on defining mouse dendritic cell populations. We thank Andrea Daniel for suggestions in experimental design. We thank Marie Iannone for running CyTOF samples. We thank Azucena Gomez-Cabrero for her assistance with designing the CyTOF panel. Graphics for schematics were created using Biorender.com. We thank Bristol Myers Squibb for providing the anti-OX40 antibody.

This work was supported by the NCI with grants 7R35CA197616 to DGK, F30CA268910 to CS, 1R38CA245204 to CLK, F30CA221268 to AJW, and P30CA14236 for the Duke Cancer Center Support Grant. Funding for this work was also provided by Varian Medical Systems, but Varian did not play a role in the design of this study or in analyzing the results.

---

**Supplementary Fig 1.**
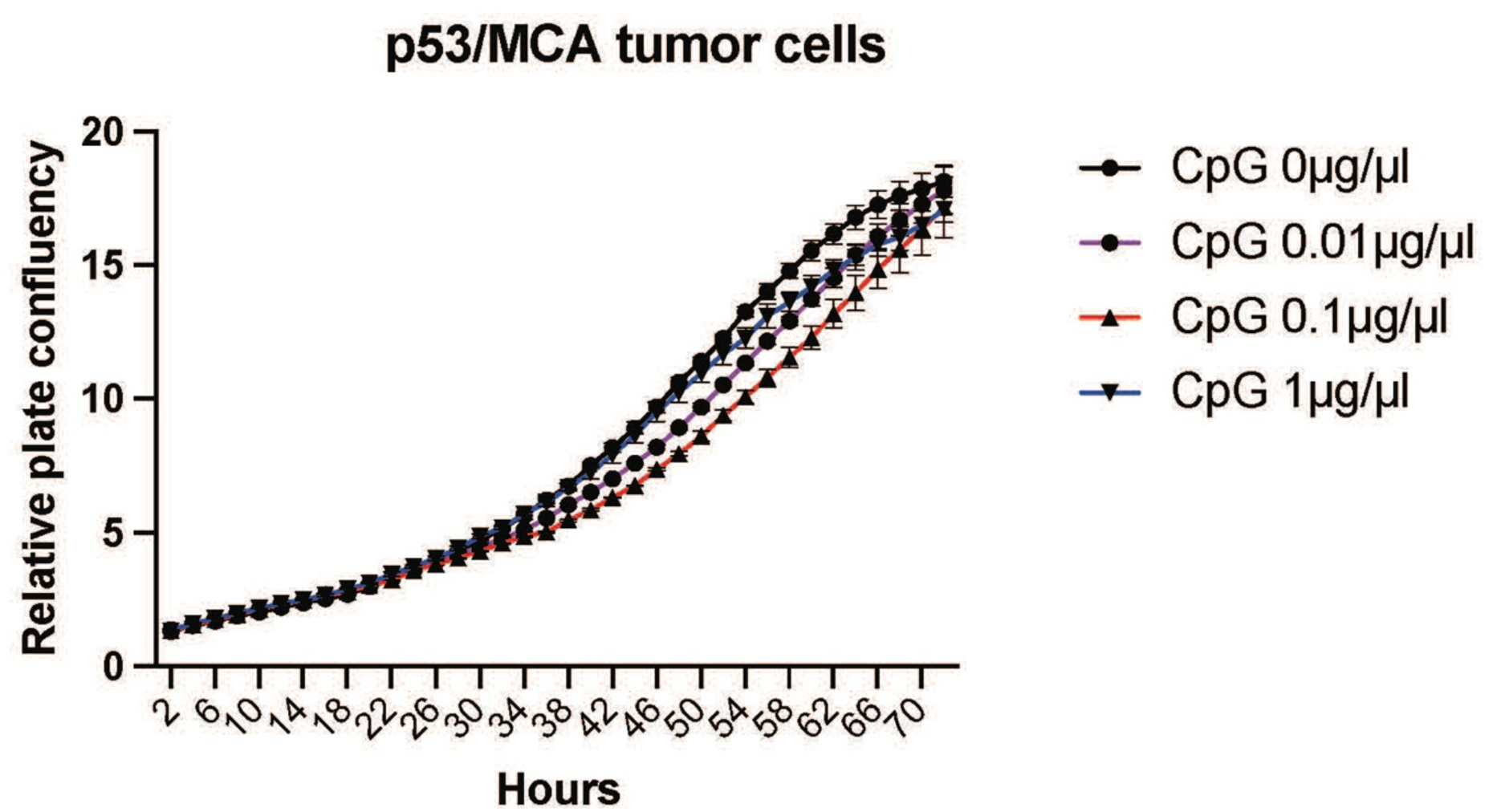
IncuCyte Live-Cell assay showed no direct effect on cell proliferation after co-incubation of p53/MCA sarcoma cells with titrated concentrations of CpG. IncuCyte cell proliferation assay of p53/MCA sarcomas incubated with four serial dilutions of CpG. The data is representative of three biological replicates.

**Supplementary Fig 2.**
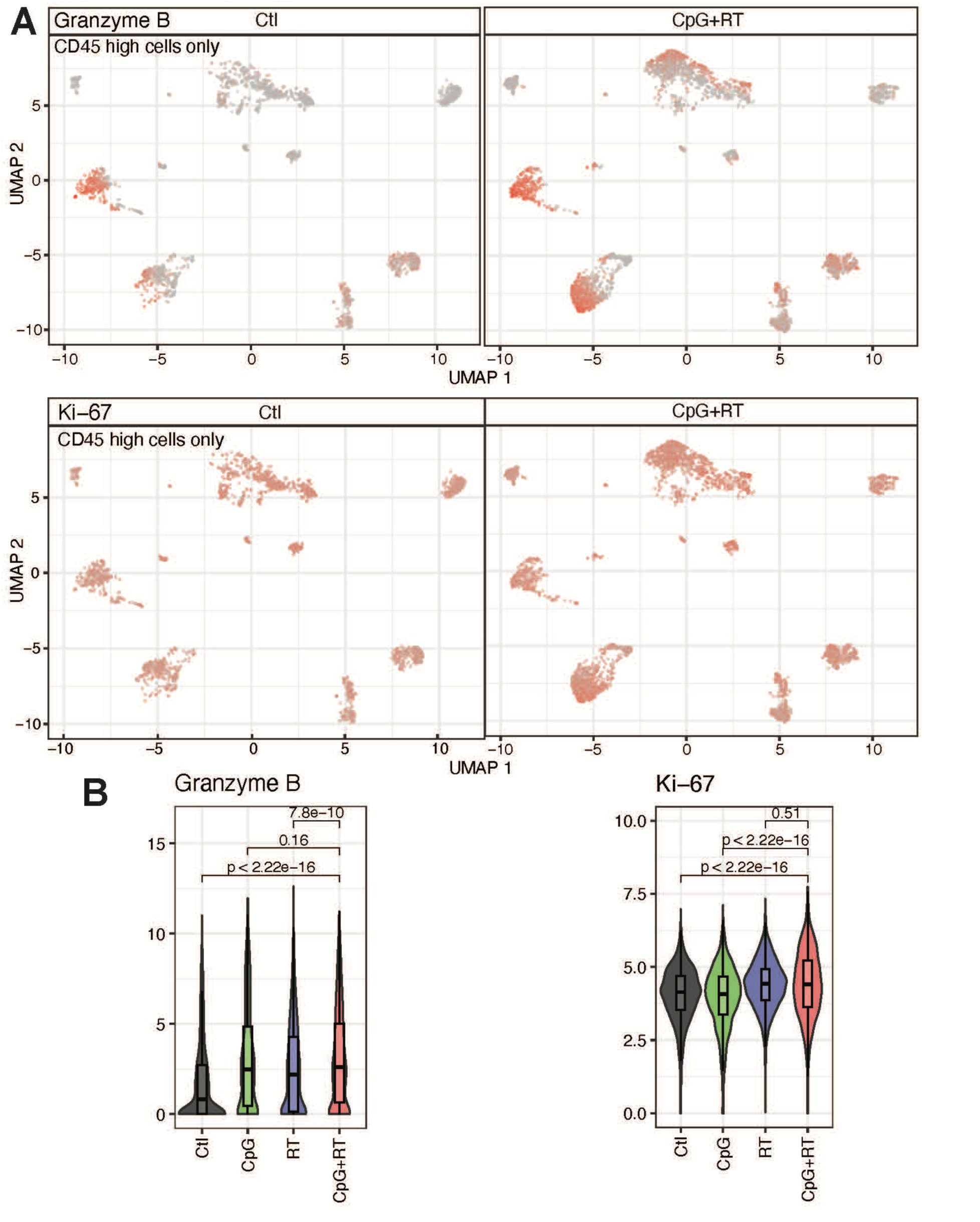
CyTOF demonstrates enhanced Granzyme B and Ki-67 expression in CD8^+^ T cells after treatment with CpG + RT. **A.** UMAP plot of CyTOF data clustering for CD45hi cells with Granzyme B and Ki67 expression colored by the ion intensity. **B.** Violin/boxplot of granzyme B and Ki-67 ion intensity in CD45 high cells across all treatment groups.

**Supplementary Fig 3.**
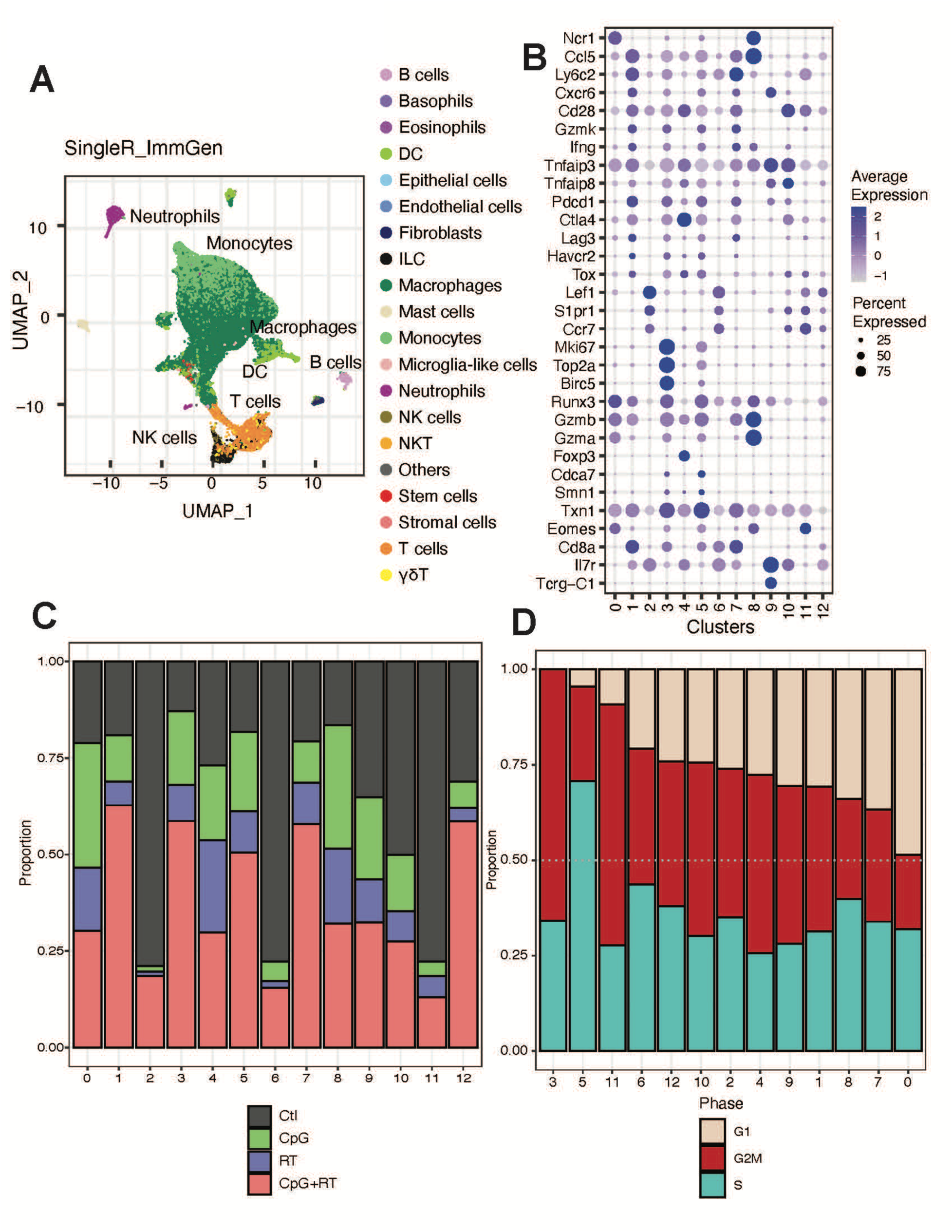
Single cell RNA-seq shows increased CD8^+^ T cells infiltration in the tumor area after CpG + RT that is highly proliferative and activated. **A.** UMAP plot of scRNA-seq clustering for all CD45hi cells from all tumors and treatment groups. **B.** Bubble plot of iteration genes in each of the T cell subclusters. **C**. Proportion of T cells contributed from each treatment group. **D**. Phases of cell cycle of clusters of different T cells and NK cells.

**Supplementary Fig 4.**
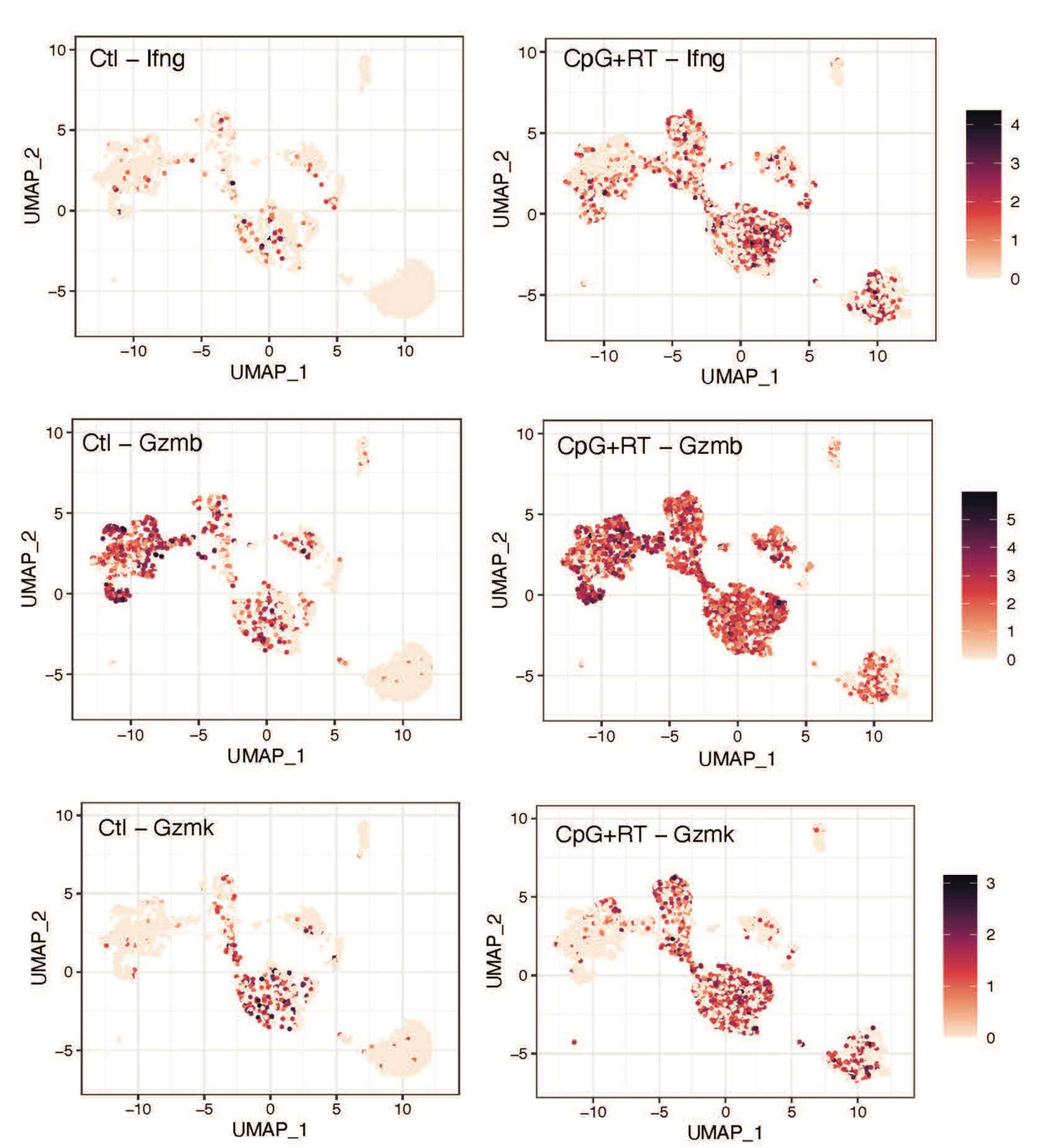
Single cell RNA-seq shows elevated expression of Granzyme B, Granzyme K, and IFNγ in CD8^+^ T cells after treatment with with CpG **+ RT.** UMAP plots of NK cells and T cells with Granzyme B, Granzyme K, and IFNγ expressing cells highlighted before and after CpG + RT treatments.

**Supplementary Fig 5.**
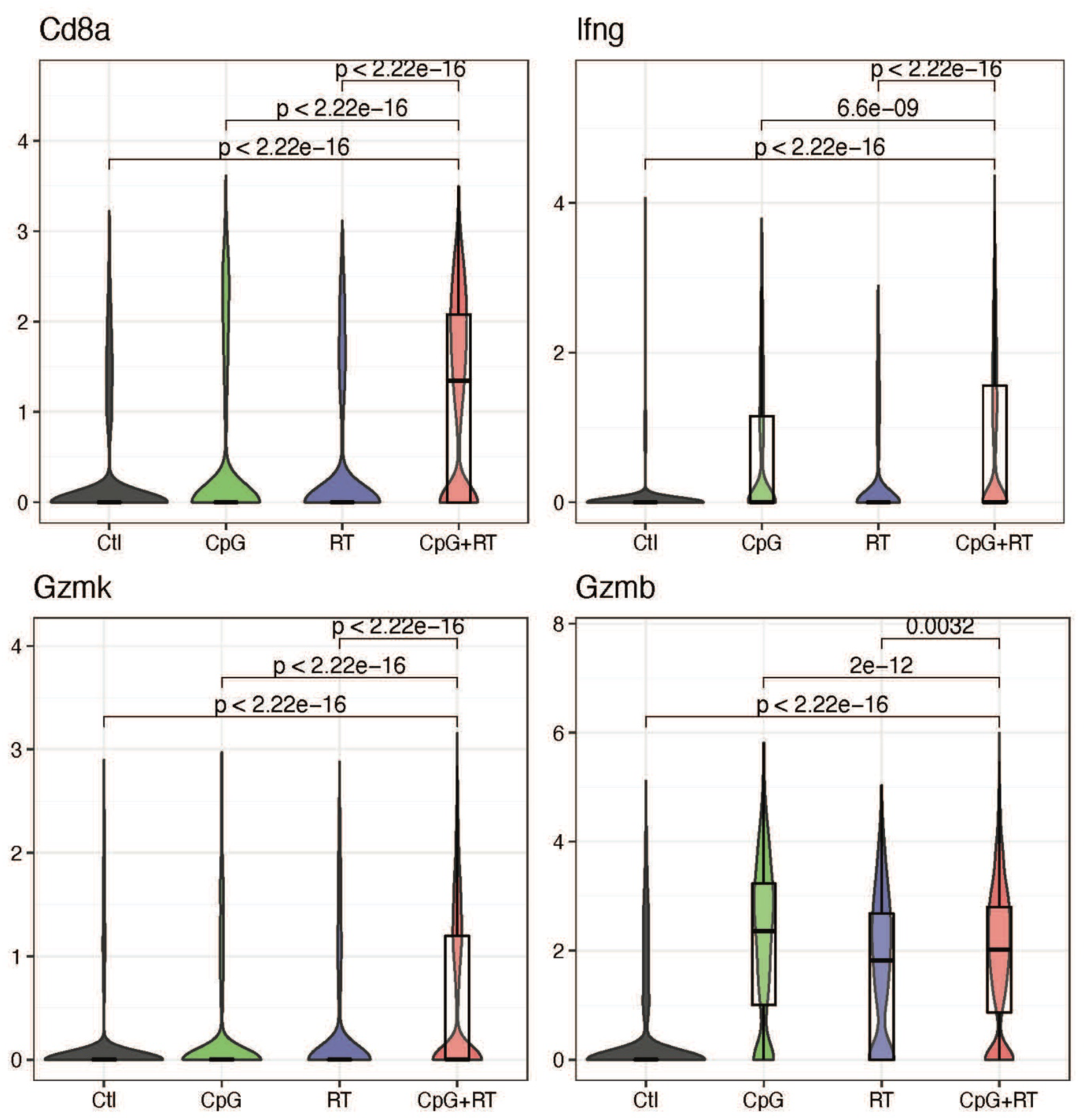
Statistical analysis and violin plots of CD8, IFNγ, Granzyme B, and Granzyme K mRNA levels in single cell RNA-seq data across different treatment groups. Normalized count of Cd8a, IFNγ, Gzmk, and Gzmb in single-cell transcriptomic data stratified by treatment group. P values were calculated using two-sided Wilcoxon tests.

**Supplementary Fig 6.**
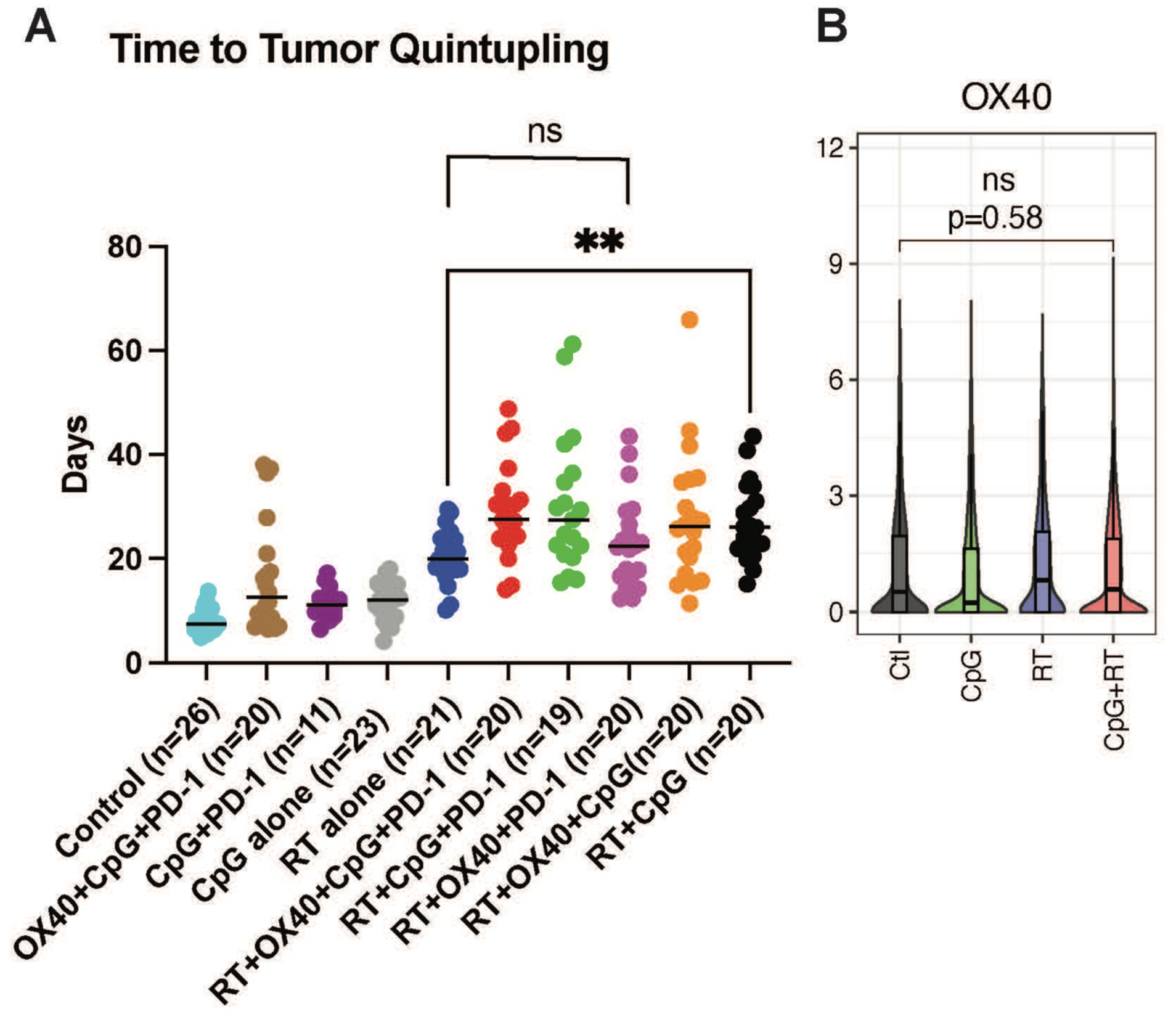
Time to tumor quintupling for p53/MCA sarcomas treated with different combination therapies. **A.** p53/MCA sarcomas were treated with anti-OX40 or vehicle control, anti-PD1 or IgG2a isotype control, CpG ODN or control GpC dinucleotides, and 0 or 20 Gy when tumors reached >70 mm^3^. Kruskal-Wallis Test was used for the group comparison, while the Wilcoxon test was selected for the pair-wise comparisons. * shows the significance of the P Value (***: p <= 0.001, ****: p <= 0.0001). **B.** Violin/boxplot of OX40 protein intensity in CD45 high cells across all treatment groups through CyTOF.

**Supplementary Fig 7.**
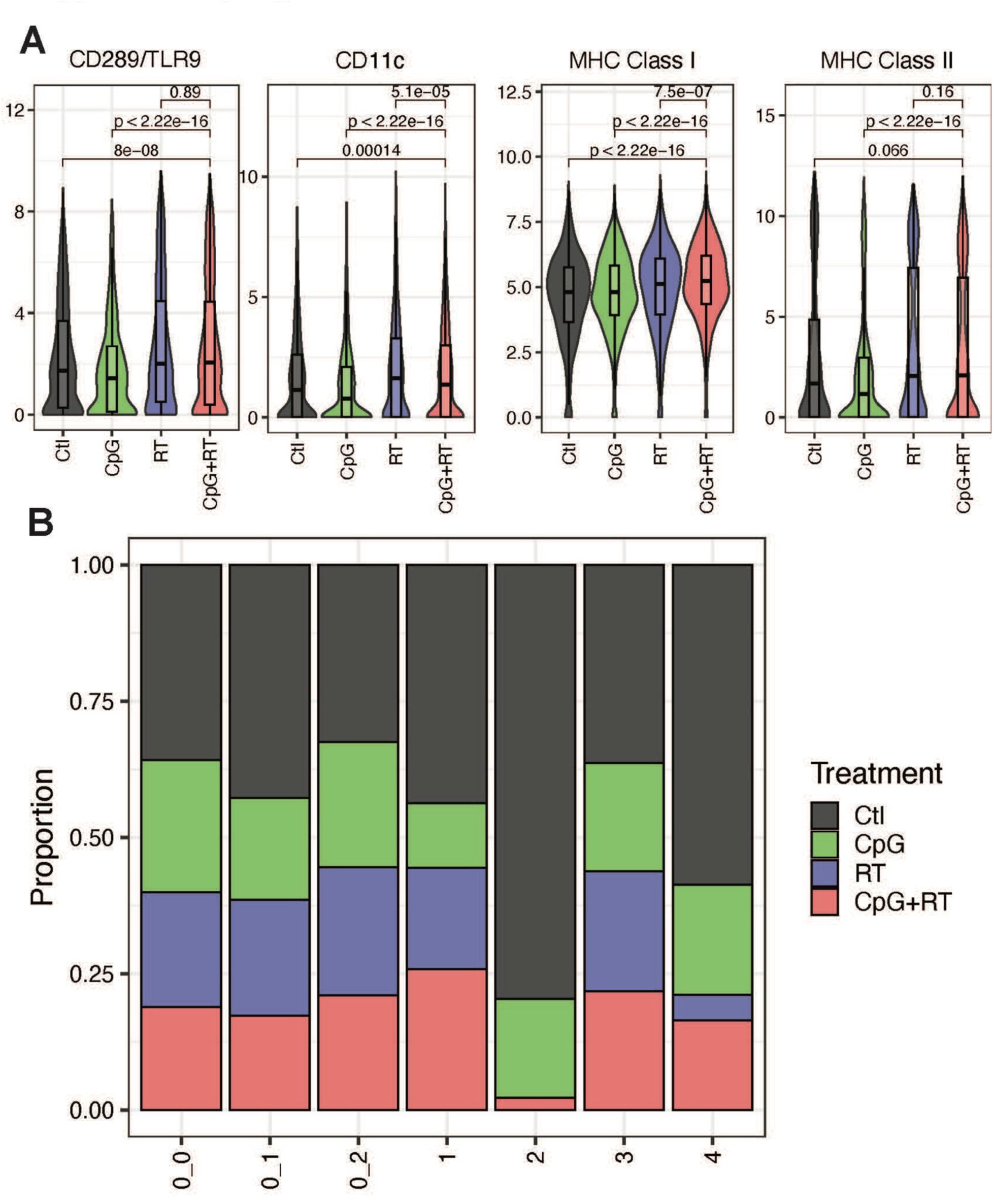
Treatment with CpG + RT promotes intratumoral dendritic cell remodeling. **A.** Violin/boxplot of CD289/TLR9, CD11c, MHC Class I, and MHC Class II protein intensity in CD45 high cells across all treatment groups through CyTOF. **B.** Proportion of DCs contributed from each treatment group.

**Supplementary Fig 8.**
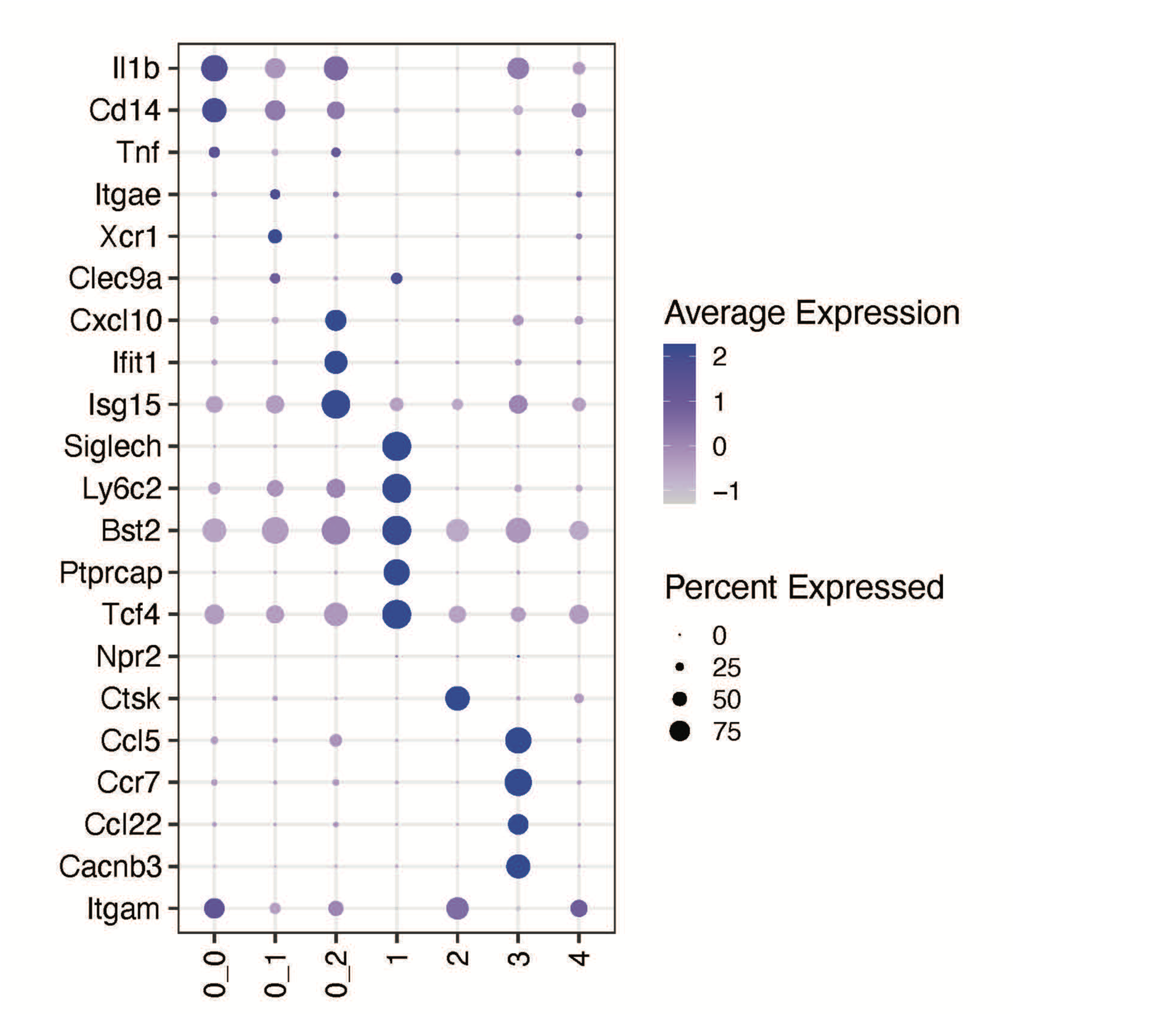
Bubble plot of iteration genes in each of the DC subclusters.

